# Improved design and analysis of practical minimizers

**DOI:** 10.1101/2020.02.07.939025

**Authors:** Hongyu Zheng, Carl Kingsford, Guillaume Marçais

## Abstract

**Motivation:** Minimizers are methods to sample *k*-mers from a sequence, with the guarantee that similar set of *k*-mers will be chosen on similar sequences. It is parameterized by the *k*-mer length *k*, a window length *w* and an order on the *k*-mers. Minimizers are used in a large number of softwares and pipelines to improve computation efficiency and decrease memory usage. Despite the method’s popularity, many theoretical questions regarding its performance remain open. The core metric for measuring performance of a minimizer is the density, which measures the sparsity of sampled *k*-mers. The theoretical optimal density for a minimizer is 1*/w*, provably not achievable in general. For given *k* and *w*, little is known about asymptotically optimal minimizers, that is minimizers with density *O*(1*/w*).

**Results:** We derive a necessary and sufficient condition for existence of asymptotically optimal minimizers. We also provide a randomized algorithm, called the Miniception, to design minimizers with the best theoretical guarantee to date on density in practical scenarios. Constructing and using the Miniception is as easy as constructing and using a random minimizer, which allows the design of efficient minimizers that scale to the values of *k* and *w* used in current bioinformatics software programs.

**Availability:** Reference implementation of the Miniception and the codes for analysis can be found at https://github.com/kingsford-group/miniception.

## 1 Introduction

The *minimizer* (Schleimer *et al.*, 2003; Roberts *et al.*, 2004a,b), also known as *winnowing*, is a method to sample positions or *k*-mers (subsequences of length *k*) from a long sequence. Given two sequences that share long enough exact subsequences, the minimizer selects the same *k*-mers in the identical subsequences, making it suitable to quickly estimate how similar two sequences are, and to quickly locate shared subsequences. The minimizers method is very versatile and is used in various ways in many bioinformatics programs (see the reviews (Rowe (2019); Marçais *et al.* (2019)) for examples) to reduce the total computation cost or the memory usage.

A minimizer has 3 parameters: (*w, k, 𝒪*). *k* is the length of the *k*-mers of interest while *w* is the length of the *window* : a least one *k*-mer in any window of *w* consecutive *k*-mers, or equivalently in any substring of length *w*+*k* −1, must be selected. Finally, is a total order (i.e., a permutation) of all the *k*-mers, and it determines how the *k*-mers are selected: in each window the minimizers selects the minimum *k*-mer according to the order 𝒪 (hence the name minimizers), and in case of multiple minimum *k*-mers, the leftmost one is selected. The main measure of performance for minimizer is the *density*: the expected number of positions selected divided by the length of the sequence. In general, minimizers with lower densities are desirable as fewer selected positions implies a further reduction in the run time or memory usage of applications using minimizers, while preserving the guarantee that for similar sequences the same *k*-mers are selected. For given parameters *w* and *k*, the choice of the order 𝒪 changes the expected density.

The density of a minimizer is lower bounded by 1*/w*, where exactly 1 position per window is selected, and upper bounded by 1, where every position is selected. A minimizer with a density of 1*/w* may not exist for every choice of *w* and *k*, and how to find an order 𝒪 with the smallest possible density for any *w* and *k* efficiently is still an open question. We are interested in constructing minimizers with density of *O*(1*/w*) — that is, within constant factor of optimal density — and avoid minimizers with density of Ω(1).

Schleimer *et al.* gave two results about the density of minimizers: under some simplifying assumptions, (1) the expected density obtained by a randomly chosen order on a random input sequence is 2*/*(*w* + 1) and (2) the density is lower-bounded by 1.5*/*(*w* + 1). Although these estimates are useful in practice, they are dependent on some hidden assumptions and do not represent the behavior of minimizers in all cases.

In previous publications we refined these results in multiple ways by looking at the asymptotic behavior of minimizers, by considering the cases where *k* is fixed and *w* ≫ *k* and where *w* is fixed and *k* ≫ *w*. First, when *w* is fixed and *k* ≫ *w*, we gave a construction of a minimizers with density of 1*/w* + *O*(*ke*^−*αk*^), for some *α* > 0 (Marçais *et al.*, 2018). That is, density arbitrarily close to the optimal 1*/w* is achievable for large values of *k*. The apparent contradiction between this result and Schleimer’s lower bound stems from a hidden assumption: *k* should not be too large compared to *w*. Second, we showed that when *k* is fixed and *w* ≫ *k*, the density of any minimizer is Ω(1). Hence, a density of 2*/*(*w* + 1), or even *O*(1*/w*), as proposed for a random order does not apply for large *w* and fixed *k* (Marçais *et al.*, 2018). In other words, this original density estimate relies on a hidden assumption: *k* should not be too small compared to *w*.

Examples of minimizers with density less than 2*/*(*w*+1) exist in practice, but these examples are one-off construction for a particular choice of parameters *w* and *k* (Marçais *et al.*, 2017). Methods to construct minimizers that have density lower than Ω(1) and work for any *w* exist: a density of 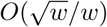 is obtained by Marçais *et al.* and Zheng *et al.* improves the result to *O*(ln(*w*)*/w*). But neither of them reaches the desired asymptotically optimal *O*(1*/w*) density. This naturally raises the question on whether a minimizer with density 2*/*(*w* + 1) or *O*(1*/w*) is possible assuming both parameters *k* and *w* can be arbitrarily large.

This paper has three main contributions. First, in Section 2.2, we prove that as *w* grows asymptotically, the condition that log_*σ*_(*w*) − *k* = *O*(1) is both a necessary and sufficient condition for the existence of minimizers with density of *O*(1*/w*). In other words, to construct asymptotically optimal minimizers, it is sufficient and necessary that the length of the *k*-mers grow at least as fast as the logarithm of the window size.

Second, in Section 2.3, by slightly strengthening the constraint on *k* —i.e., *k* ≥ (3 + *ϵ*) log_*σ*_(*w*), for any fixed *ϵ* > 0— we show that a random minimizer has expected density of 2*/*(*w* + 1) + *o*(1*/w*). This theorem is a direct extension of the result by Schleimer *et al.* as it removes any hidden assumptions and gives a sufficient condition for the result to hold.

Third, in Section 2.4, we give a construction of minimizers, called the *Miniception*, with expected density on a random sequence of 1.67*/w* + *o*(1*/w*) when *k* ≈ *w*. This is an example of minimizers with guaranteed density < 2*/*(*w* + 1) that works for infinitely many *w* and *k*, not just an ad-hoc example working for one or a small number of parameters *w* and *k*. This is also the first example of a family of minimizers with guaranteed expected density < 2*/*(*w* + 1) that works when *k* ≈ *w* instead of the less practical case of *k* ≫ *w*. Moreover, unlike other methods with low density in practice (Orenstein *et al.*, 2016; Ekim *et al.*, 2020; DeBlasio *et al.*, 2019), the Miniception does not require the use of expensive heuristics to precompute and store a large set of *k*-mers. Selecting *k*-mers with the Miniception is as efficient as a selecting *k*-mers with a random minimizer using a hash function, and does not require any additional storage.

In Section 2 we give detailed proof of our results, in Section 3 we compare the Miniception to existing methods to create minimizers, and in section 4, we discuss further possible directions for the theory and practical applications of minimizers.

## 2 Methods

### 2.1 Preliminary

In this section, we restate several theorems from existing literature that are useful for later sections. In the following, Σ = {0, 1, …, *σ* − 1} is an alphabet (mapped to integers) of size *σ* and we assume that *σ* ≥ 2 and is fixed. If *S* ∈ Σ* is a string, then |*S*| denotes the length of *S*.

#### Definition 1 (Minimizer and Windows).

*A “minimizer” is characterized by* (*w, k*, 𝒪) *where w and k are integers and* O *is a complete order of* Σ^*k*^. *A “window” is a sequence of length* (*w* + *k* − 1), *consisting of exactly w over-lapping k-mers. Given a window as input, the minimizer outputs the location of the smallest k-mer according to, breaking ties by preferring leftmost k-mer.*

When 𝒪 is the dictionary order, it is called a *lexicographic minimizer*. A minimizer created by randomly choosing a permutation of Σ^*k*^ uniformly over all possible permutations is called a *random minimizer*. See Figure 1 for an illustration of these and the following concepts.

**Figure 1:**
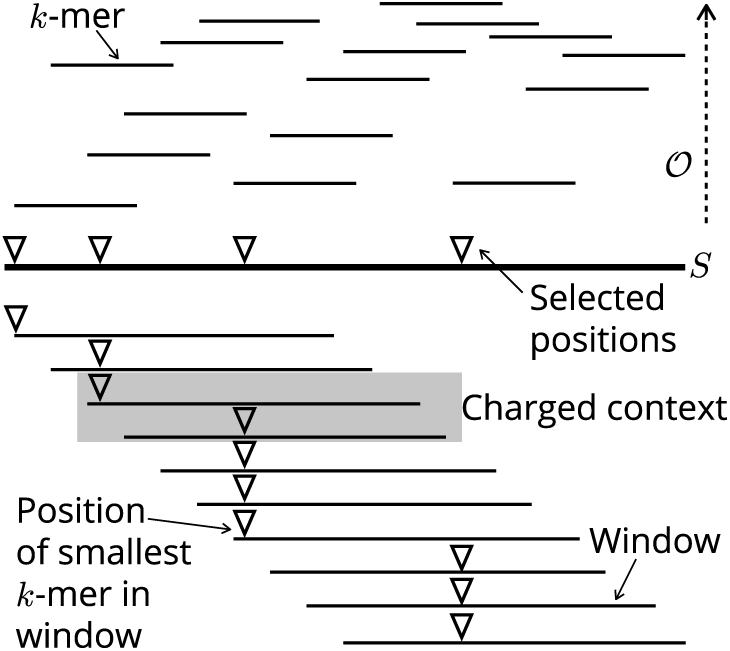
The sequence S is broken into k-mers. In each window (w consecutive k-mers) the position of the lowest (smallest according to 𝒪) k-mer is selected. The gray context (2 consecutive windows) is an example of a charged because the first and second window selected different positions. There are a total of 3 charged contexts in this example.

#### Definition 2 (Density).

*Given a sequence S* ∈ Σ* *and a minimizer, a position in S is selected if the minimizer picks the k-mer at that position in any window of w consecutive k-mers. The specific density of a minimizer on S is the number of selected positions divided by the total number of k-mers in S. The density of a minimizer is the expected specific density on a sufficiently long random string.*

Note that the density is calculated by expectation over random sequences, and is independent from *S*.

#### Definition 3 (Charged Contexts).

*A “context” of S is a substring of length* (*w* + *k*), *or equivalently*, 2 *overlapping windows. The minimizer is applied to both the first and last windows, and a context is “charged” if different positions are picked.*

#### Definition 4 (Density Factor).

*The density factor of a minimizer is its density multiplied by* (*w* + 1). *Intuitively, this is the expected number of selected locations in a random context.*

We denote by *W*_*w,k*_ = Σ^*w*+*k*−1^ the set of all possible windows and *C*_*w,k*_ = Σ^*w*+*k*^ the set of all contexts. We may drop the subscript *w, k* when these parameters are clear from the context.

For minimizers, the definition of charged contexts can be simplified. The following lemma can be found in previous literature (Schleimer *et al.* (2003)). We prove it here for clarity.

#### Lemma 1 (Charged Contexts of Minimizers).

*For a minimizer, a context is charged if and only if the minimizer picks either the first k-mer of the first window or last k-mer in the last window.*

*Proof.* On one hand, if the minimizer picks either the first or the last *k*-mer in the context, it cannot be picked in both windows. On the other hand, if the minimizer does not pick either the first or the last *k*-mer, it will pick the same *k*-mer in both windows. Assuming otherwise, both picked *k*-mers is in both window so this means one of them is not minimal, leading to a contradiction.

Intuitively, as a context is two overlapping windows, a charged context corresponds to the event that a new *k*-mer is picked in the latter window. Thus, counting picked *k*-mers is equivalent to counting charged contexts.

#### Lemma 2 (Density by Random Contexts).

*The density of a minimizer equals the probability that a context drawn from uniform distribution over* Σ^*w*+*k*^ *is charged, or equivalently, is equal to the specific density over a circular de Bruijn sequence of order w* + *k (the circular string of length* Σ^*w*+*k*^ *that contains each* (*w*+*k*)*-mer exactly once).*

A related concept is *universal hitting sets* (UHS) (Orenstein *et al.*, 2016), which is central to the analysis of minimizers (Marçais *et al.*, 2017, 2018; Ekim *et al.*, 2020).

#### Definition 5 (Universal Hitting Sets).

*Assume U* ⊆ Σ^*k*^. *If U intersects with every w consecutive k-mers (or equivalently, the set of k-mers in every S* ∈ *W*_*w,k*_*), it is a UHS over k-mers with path length w and relative size* |*U* |*/σ*^*k*^.

#### Lemma 3 (Minimal Decycling Sets).

*(Mykkeltveit (1972)) Any UHS over k-mers has relative size at least* 1*/k* − *o*(1*/k*).

### 2.2 Condition for Asymptotically Optimal Minimizers

In this section, culminating with Theorem 2, we prove that to have asymptotically optimal minimizers (i.e., minimizers with density of *O*(1*/w*)), the length *k* of the *k*-mers must be sufficiently large compared to the length *w* of the windows, and that this condition is necessary and sufficient. To be more precise, we treat *k* as a function of *w*, and study the density as *w* grows to infinity. We show that the lexicographic minimizers is asymptotically optimal provided that *k* is large enough: log_*σ*_(*w*) − *k* = *O*(1). This result may be surprising as in practice the lexicographic minimizers has high density (Roberts *et al.* (2004a); Marçais *et al.* (2017) and see Section 4). One interpretation of Theorem 2 is that asymptotically, all minimizers behave the same regarding density.

#### 2.2.1 Minimizers with exceedingly small *k*

If *k* is exceedingly small, in the sense that *k* does not even grow as fast as the logarithm of *w* —i.e., log_*σ*_(*w*) − *k* → ∞ as *w* grows— no minimizer will obtain density *O*(1*/w*). To see this, for any order 𝒪, let *y*_O_ be the smallest of all *k*-mers. Any context starting with *y*_O_ is charged, and the proportion of such context is *σ*^−*k*^. The density calculated from these contexts only is already > Θ(1/*w*), as 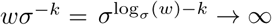.

For this reason, in the following we are only interested in the case where *k* is large enough. That is, there exists a fixed constant *c* such that *k* ≥ log_*σ*_(*w*) − *c* for all sufficiently large *w*.

#### 2.2.2 Lexicographic Minimizers

We first prove the special case that the lexicographic minimizer achieves *O*(1*/w*) density with parameter *k* = ⌊log_*σ*_ (*w/*2)⌋ − 2. Recall *W*_*k,w*_ = Σ^*k*+*w*−1^ is the set of all windows, and *C*_*k,w*_ = Σ^*k*+*w*^ is the set of all contexts. Let *z* ∈ *W*_*k,w*_ be a window, *f* (*z*) : *W*_*k,w*_ → {0, 1, …, *w* − 1} be the minimizer function, and 𝒞 ⊂ *C*_*k,w*_ be the set of charged contexts for this minimizer. Let *W*^+^ = {*z* ∈ *W* | *f*(*z*) = 0}, the set of windows where the minimizer picks the first *k*-mer, and similarly *W*^−^ = {*z* ∈ *W* | *f*(*z*) = *w* − 1}. This is a corollary to Lemma 1:

##### Corollary 1.

|𝒞| ≤ *σ*(|*W*^+^| + |*W*^−^|).

We now use the notion # to denote any nonzero character of Σ, and 0^*d*^ to denote *d* consecutive zeroes. Let *A*′^+^ be the set of windows whose first *k*-mer is 0^*k*^, and for 1 ≤ *i* ≤ *k*, let 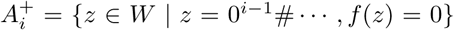, that is, the set of windows that starts with exactly *i* − 1 zeros and have the minimizer function pick the first *k*-mer. All 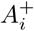 and *A*^+^ are mutually disjoint. Since the minimizer always pick 0^*k*^ at the start of the window, we have 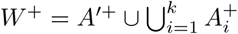.

##### Lemma 4.

*For* 1 ≤ *i* ≤ *k*, 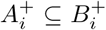, *where* 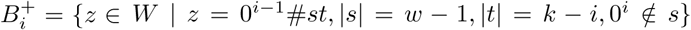, *that is, the set of windows that starts with* 0^*i*−1^# *and does not contain* 0^*i*^ *in the next w* − 1 *bases.*

*Proof.* We need to show that if a window *z* starts with 0^*i*−1^# and is not in 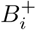, *f*(*z*) ≠ 0. As 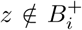, there is a stretch of 0^*i*^ in *z* before the last *k* − *i* characters. This means there is a *k*-mer of form 0^*i*^ … in *z*, and since the first *k*-mer is of form 0^*i*−1^# …, the minimizer will never pick the first *k*-mer.

In our previous paper (Zheng *et al.* (2020)), we proved that

##### Lemma 5.

*The probability that a random string of length l does not contain* 0^*d*^ *anywhere is at most* 3(1 − 1*/σ*^*d*+1^)^*l*^.

Setting *l* = *w* − 1 and *d* = *i* and noting there are *σ*^*k*−*i*^ choices for *t* in *B*_*i*_, we have the following corollary:

##### Corollary 2.

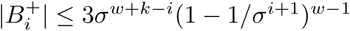.

Let *b*_*i*_ = 3*σ*^*w*+*k*−*i*^(1 − 1*/σ*^*i*+1^)^*w*−1^. Combining Corollary 2 with the fact that |*A*′^+^| = *σ*^*w*−1^, we have:

##### Corollary 3.

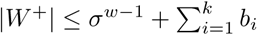.

We bound 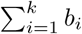 by showing that *b*_*i*_ grows exponentially with *i*.

##### Lemma 6.

*b*_*i*_ > 2*b*_*i*−1_ *for* 2 ≤ *i* ≤ *k.*

*Proof.* We have the following:

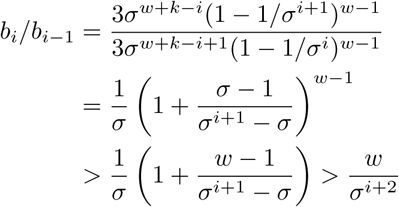

Note that we also use the fact (1 + *x*)^*t*^ > 1 + *xt* and *σ* − 1 ≥ 1 in the last line. The right-hand side is minimum when *i* = *k*. By our choice of *k, σ*^*k*+2^ < *w/*2, so the term is lower bounded by 2.

##### Corollary 4.

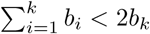

Since *b*_*k*_ = *O*(*σ*^*w*^), combining the fact |*A*′^+^| = *σ*^*w*−1^ with Corollary 3 and 4, we have |*W*^+^| = *O*(*σ*^*w*^).

The bound for |*W*^−^| is computed similarly. It is different from *W*^+^ as in case of ties for the minimal *k*-mer, the leftmost one is picked. Hence, we define *W*\* as the set of windows such that the last *k*-mer is one of the minimal *k*-mers in the window. We have *W*^−^ ⊆ *W*\*, as the last *k*-mer needs to be the minimal, with no ties, for the minimizer to pick it.

Similarly, we define *A*′^−^ as the set of windows that ends with 0^*k*^, and for 1 ≤ *i* ≤ *k*, define 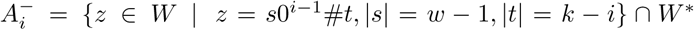 that is, the set of windows whose last *k*-mer starts with 0^*k*−1^# while satisfying the condition for *W*\*. Again, *A*′^−^ and all 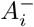 are mutually disjoint, and we have 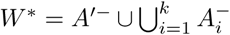. There is an analogous lemma for bounding 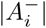:

##### Lemma 7.

*For* 1 ≤ *i* ≤ *k*, 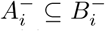, *where* 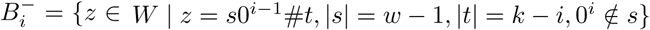.

*Proof.* We need to show that if a window ends with a *k*-mer of form 0^*i*−1^#*t* and contains 0^*i*^ before last *k*-mer, it is not in *W*\*. In that case the window contains a *k*-mer of form 0^*i*^ …, which is strictly smaller than the last *k*-mer of the form 0^*i*−1^# …, violating the condition of *W*\*.

Note that 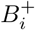 and 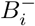 have highly similar expressions. In fact, we can simply bound the size of 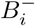 by *b*_*i*_ using the identical argument. This immediately means we have the exactly same bound for |*W*\*| and |*W*^+^|, as *A*′^−^ also has the same size as *A*′^+^.

##### Theorem 1.

*The lexicographic minimizer with k*_0_ = ⌊log_*σ*_ (*w/*2)⌋ − 2 *has density O*(1*/w*).

*Proof.* We have | 𝒞| ≤ *σ*(|*W*^+^| + |*W*^−^|) ≤ *σ*(|*W*^+^| + |*W*\*|) = *O*(*σ*^*w*^), and the density is 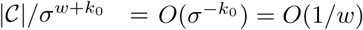.

Next, we extend this result to show that this bound holds for all *k* as long as *k* > log_*σ*_(*w*)−*c* for some constant *c*. As *k*_0_ < log_*σ*_ *w*, the following lemmas establishes our claim for small and large *k*:

##### Lemma 8.

*The lexicographic minimizer with k* = *k*_0_ − *c for constant c* ≥ 0 *has density O*(1*/w*).

##### Lemma 9.

*The lexicographic minimizer with k* > *k*_0_ *has density O*(1*/w*).

We prove both lemmas in Section S1. Combining everything we know, we have the following theorem.

##### Theorem 2.

*For k* ≥ log_*σ*_(*w*) − *c with constant c, the lexicographic minimizer achieves density of O*(1*/w*). *Otherwise, no minimizer can achieve density of O*(1*/w*).

### 2.3 Density of Random Minimizers

We now study the density of random minimizers. Random minimizers are of practical and theoretical interest. In practice, implementing a random minimizer is relatively easy using a hash function, and these minimizers usually have lower density than lexicographic minimizers. Consequently, most software programs using minimizers use a random minimizer. The constant hidden in the big-*O* notation of Theorem 2 may also be too large for practical use, while later in Theorem 3, we guarantee a density of 2*/*(*w* + 1) with a slightly more strict constraint over *k*.

Schleimer *et al.* (2003) estimated the expected density of random minimizers to be 2*/*(*w* + 1) with several assumptions on the sequence (which do not strictly hold in practice), and our main theorem (Theorem 3) achieves the same result up to *o*(1*/w*) with a single explicit hypothesis between *w* and *k*. Combined with our previous results on connecting UHS to minimizers, we also provide an efficient randomized algorithm to construct compact universal hitting sets with 2 + *o*(1) approximation ratio.

#### 2.3.1 Random minimizers

In the estimation of the expected density for random minimizers, there are 2 sources of randomness: (1) the order 𝒪 on the *k*-mers is selected at random among all the permutations of Σ^*k*^ and (2) the input sequence is a very long random string with each character chosen IID. The key tool to this part is the following lemma to control the number of “bad cases” when a window contains two or more identical *k*-mers. Chikhi *et al.* (2015a) proved a similar statement with slightly different methods.

##### Lemma 10.

*For any ϵ* > 0, *if k* > (3 + *ϵ*) log_*σ*_ *w, the probability that a random window of w k-mers contains two identical k-mers is o*(1*/w*).

*Proof.* We start with deriving the probability that two *k*-mers in fixed locations *i* and *j* are identical in a random window. Without loss of generality, we assume *i* < *j*. If *j* − *i* ≥ *k*, the two *k*-mers do not share bases, so given they are both random *k*-mers independent of each other, the probability is *σ*^−*k*^ = 1*/w*^3+ *ϵ*^ = *o*(1*/w*^3^).

Otherwise, the two *k*-mers intersect. We let *d* = *j* − *i*, and *m*_*i*_ to denote *i*^th^ *k*-mer of the window. We use *x* to denote the subsequence from the start of *m*_*i*_ to the end of *m*_*j*_ with length *k* + *d* (or equivalently, the union of *m*_*i*_ and *m*_*j*_). If *m*_*i*_ = *m*_*j*_, the *n*^th^ character of *m*_*i*_ is equal to the *n*^th^ character of *m*_*j*_, meaning *x*_*n*_ = *x*_*n*+*d*_ for all 0 ≤ *n* < *k*. This further means *x* is a repeating sequence of period *d*, so *x* is uniquely determined by its first *d* characters and there are *σ*^*d*^ possible configurations of *x*. The probability a random *x* satisfies *m*_*i*_ = *m*_*j*_ is then *σ*^*d*^*/σ*^*k*+*d*^ = *σ*^−*k*^ = *o*(1*/w*^3^), which is also the probability of *m*_*i*_ = *m*_*j*_ for a random window.

The event that the window contains two identical *k*-mers is the union of events of form *m*_*i*_ = *m*_*j*_ for *i* < *j*, and each of these events happens with probability *o*(1*/w*^3^). Since there are Θ(*w*^2^) events, by the union bound, the probability that any of them happens is upper bounded by *o*(1*/w*).

We are now ready to prove the main theorem of this section.

##### Theorem 3.

*For k* > (3 + *ϵ*) log_*σ*_(*w* + 1), *the expected density of a random minimizer is* 2*/*(*w* + 1) + *o*(1*/w*).

*Proof.* Given a context *c* ∈ Σ^*w*+*k*^, we use *I*(*c*) to denote the event that *c* has two identical *k*-mers. As *c* has (*w* +1) *k*-mers, by Lemma 10, *P*_*c*_(*I*(*c*)) = *o*(1*/w*) assuming *c* is a random context.

Recall that a random minimizer means the order 𝒪 is randomized, and 𝒞 is the set of charged contexts. For any context *c* that does not have duplicate *k*-mers, we claim *P*_𝒪_ (*c* ∈ 𝒞) = 2*/*(*w* + 1). This is because given all *k*-mers in *c* are distinct, under the randomness of, 𝒪 each *k*-mer has probability of exactly 1*/*(*w* + 1) to be the minimal. By Lemma 1, *c* ∈ 𝒞 if and only if the first or the last *k*-mer is the minimal, and as these two events are mutually exclusive, the probability of either happening is 2*/*(*w* + 1). The expected density of the random minimizer then follows:

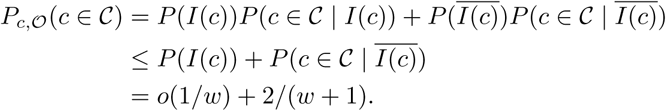

#### 2.3.2 Approximately Optimal Universal Hitting Sets

One interesting implication of Theorem 3 is on construction and approximation of compact universal hitting sets. In our previous paper (Zheng *et al.* (2020)), we proved a connection between universal hitting sets and forward schemes. We restate it with minimizers, as follows.

##### Theorem 4.

*For any minimizer* (*w, k*, 𝒪), *the set of charged contexts over a de Bruijn sequence of order w*+*k is a UHS over* (*w* + *k*)*-mers with path length w, and with relative size identical to the density of the minimizer.*

We discuss the UHS with *w* > *k* and *w* < *k* separately. In the first case, the best UHS has its relative size lower bounded by the decycling set, and in the second case, a stronger bound of 1*/w* is available.

##### Lemma 11.

*For sufficiently large k and w* > *k* − (3 + *ϵ*) log_*σ*_ (*k*+1), *there exists a universal hitting set of relative size* 2*/k* + *o*(1*/k*) *with path length w. This is a* 2 + *o*(1) *approximation of the minimum size universal hitting set.*

*Proof.* We pick *k*′ = (3 + *ϵ*) log_*σ*_ (*k* + 1) (ignoring rounding for simplicity), and let *w*′ = *k* − *k*′. By our setup, *w* ≥ *w*′, and *k*′ = *o*(*k*) (implying *w*′ = *k* − *o*(*k*)). By Theorem 3, the charged context set of a random minimizer (*w*′, *k*′, *𝒪*′) has relative size 2*/w*′ + *o*(1*/w*′) = 2*/k* + *o*(1*/k*). By Theorem 4, the charged context set is a UHS over *k*-mers with path length *w*. This finishes the first part of the proof.

For the second part, by Lemma 3 the minimum size UHS, regardless of path length, will have relative size at least 1*/k* − *o*(1*/k*). As our proposed UHS has relative size 2*/k* + *o*(1*/k*), it is a 2 + *o*(1) approximation of the minimum size UHS.

##### Lemma 12.

*For sufficiently large k and w* ≤ *k* − (3 + *ϵ*) log_*σ*_ (*k*+1), *there exists a universal hitting set of relative size* 2*/w* + *o*(1*/w*) *with path length w. This is a* 2 + *o*(1) *approximation of the best possible universal hitting set.*

*Proof.* We pick *w*′ = *w* and *k*′ = *k* − *w*. By our setup, *k*′ ≥ (3 + *ϵ*) log_*σ*_(*w*′ + 1) as *w* < *k*. By the identical argument as seen in the last lemma, we obtain a UHS over *k*-mers with path length *w* and relative size 2*/w* +*o*(1*/w*).

For the second part, note that the minimal relative size of a UHS with path length *w* is 1*/w* as it need to hit every one of every *w k*-mer on the de Bruijn sequence of order *k*, where every *k*-mer appears exactly once. As our set has relative size 2*/w* + *o*(1*/w*), it is a 2 + *o*(1) approximation.

These two lemmas give us the desired result. We say an algorithm for constructing UHS is efficient if it runs in poly(*w, k*)*σ*^*w*+*k*^ time, as the output length of such algorithms is already at least *σ*^*w*+*k*^.

##### Theorem 5.

*For sufficiently large k and for arbitrary w, there exists an efficient randomized algorithm to generate a* (2 + *o*(1))*-approximation of a minimum size universal hitting set over k-mers with path length w.*

This is also the first known efficient algorithm to achieve constant approximation ratio, as previous algorithms (Orenstein *et al.*, 2016; Ekim *et al.*, 2020; DeBlasio *et al.*, 2019) use path cover heuristics with approximation ratio dependent on *w* and *k*.

### 2.4 The Miniception

In this section, we develop a minimizer (or rather, a distribution of minimizers) with expected density strictly below 2*/w* + *o*(1*/w*). The construction works as long as *w* < *xk* for some constant *x*. One other method exists to create minimizers with density below 2*/w* (Marçais *et al.*, 2018), but it requires *w* ≪ *k*, a much more restrictive condition.

The name “Miniception” is shorthand for “Minimizer Inception”, a reference to its construction that uses a smaller minimizer to construct a larger minimizer. In the estimation of the expected density of Miniception minimizers, there are multiple sources of randomness: the choice of orders in the small and in the large minimizers, and the chosen context. The construction and the proof of the Miniception uses these sources of randomness to ensure its good performance on average.

#### 2.4.1 A Tale of two UHSes

Universal hitting sets are connected to minimizers in two ways. The first connection, via the charged context set of a minimizer, is described in Theorem 4. The second and more known connection is via the idea of *compatible minimizers*. Detailed proof of the following properties are available in Marçais *et al.* (2017, 2018).

##### Definition 6 (Compatibility).

*A minimizer* (*w, k, 𝒪*) *is said to be compatible with a universal hitting set U, if the path length of U is at most w and for any m* ∈ *U, m*′ ∉ *U, m* < *m*′ *under 𝒪.*

##### Lemma 13 (Properties of Compatible Minimizers).

*If a minimizer is compatible with a UHS, (1) any k-mer outside the UHS will never be picked by the minimizer, and (2) the relative size of the UHS is an upper bound to the density of the minimizer.*

##### Theorem 6.

*Any minimizer* (*w, k, 𝒪*) *is compatible with the set of selected k-mers over a de Bruijn sequence of order w* + *k.*

The Miniception is a way of constructing minimizers that uses the universal hitting sets in both ways. Assume we have a minimizer (*w*_0_, *k*_0_, *𝒪*_0_). By Theorem 4, its charged context set 𝒞_0_ is a UHS over (*w*_0_ + *k*_0_)-mers with path length *w*_0_. According to Definition 6 and Theorem 6, we can construct a minimizer (*w, k, 𝒪*) that is compatible with 𝒞_0_, where *k* = *w*_0_ + *k*_0_, *w* ≥ *w*_0_ and any *k*-mer in 𝒞_0_ is less than any *k*-mer outside 𝒞_0_ according to 𝒪.

We assume that the smaller minimizer (*w*_0_, *k*_0_, 𝒪_0_) is a random minimizer, and that the larger minimizer (*w, k*, 𝒪) is a random compatible minimizer (meaning the order of *k*-mers within 𝒞_0_ is random in 𝒪). The Miniception is formally defined as follows:

##### Definition 7 (The Miniception).

*Given parameters w, k and k*_0_, *set w*_0_ = *k* − *k*_0_. *The Miniception is a minimizer with parameters w and k constructed as follows:*

- *A random minimizer* (*w*_0_, *k*_0_, 𝒪_0_) *called the “seed” is generated.*
- *The set of charged contexts* 𝒞_0_ ⊂ Σ^*k*^ *is calculated from the seed minimizer (note that k* = *w*_0_ + *k*_0_).
- *the order* 𝒪 *of the resulting minimizer is constructed by generating a random order within* 𝒞_0_, *and having every other k-mer compare larger than any k-mers in* 𝒞_0_.

Note that by Lemma 13, the order within *k*-mers outside 𝒞_0_ doesn’t matter in constructing 𝒪. In the following three sections, we will prove the following theorem:

##### Theorem 7.

*With w* = *w*_0_ +1, *k* = *w*_0_ +*k*_0_ *and k*_0_ > (3+ *ϵ*) log_*σ*_ (2*w*_0_ + 2), *the expected density of the Miniception is upper bounded by* 1.67*/w* + *o*(1*/w*).

As *k* = *w*_0_ + *k*_0_, for large values of *w*_0_, we can take for example *k*_0_ = 4 log_*σ*_ *w*_0_, meaning *w* ≈ *k* in these cases. This makes the Miniception the only known construction with guaranteed density < 2*/*(*w* + 1) + *o*(1*/w*) and with practical parameters.

Figure 2 provides an example of the Miniception. The Miniception can be implemented efficiently in practice. Assuming that a random order is computed with a hash function in *O*(*k*) time for a *k*-mer, determining the set of picked *k*-mers in a sequence *S* in takes *O*(*k* |*S*|) time. This is as fast as a random minimizer. In particular, there is no need to precompute the set 𝒞_0_. We discuss the implementation in more detail in Section S3, and provide a reference implementation in the GitHub repository.

**Figure 2:**
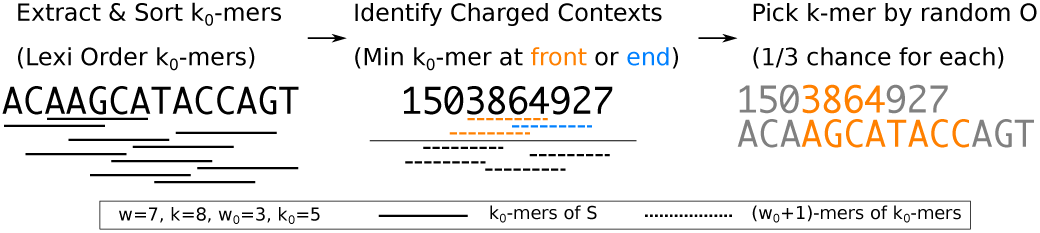
An example of running the Miniception in a window. The k_0_-mers and (w_0_ + 1)-mers are displayed by their order in 𝒪 and 𝒪_0_, where the minimal elements appear at the top. We take 𝒪_0_ to be lexicographic order for simplicity, and 𝒪 is a random order. The idea of sorting k_0_-mers will be important in deriving the theoretical guarantees of the Miniception.

#### 2.4.2 The Permutation Argument

We now focus on the setup outlined in Theorem 7. Our goal is to measure the density of the Miniception, which is equivalent by Lemma 2 to measuring the expected portion of charged contexts, that is, *P* (*c* ∈ 𝒞). There are three sources of randomness here: (1) the randomness of the seed minimizer, (2) the randomness of the order within 𝒞_0_ (which we will refer to as the randomness of 𝒪), and (3) the randomness of the context.

A context of the Miniception is a (*w* + *k*) = (2*w*_0_ + *k*_0_ + 1)-mer, which contains (2*w*_0_ + 2) *k*_0_-mers. By our choice of *k*_0_ and Lemma 10, the probability that the context contains two identical *k*_0_-mers is *o*(1*/w*_0_) = *o*(1*/w*) (as *w* = *w*_0_ + 1). Similar to our reasoning in proving Theorem 3, let *I*_0_ denote the event of duplicate *k*_0_-mers in a Miniception context:

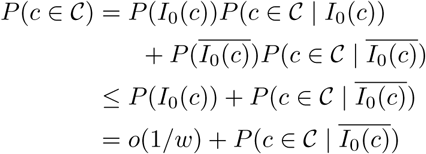

We now consider a fixed context that has no duplicate *k*_0_-mers. Recall the way we determine whether a *k*-mer is in 𝒞_0_: we check if it is a charged context of the seed minimizer, which involves only comparisons between its constituent *k*_0_-mers. This means given a context of the Miniception, we can determine if each of its *k*-mer is in the UHS 𝒞_0_ only using the order between all *k*_0_-mers. We use Ord(*c*) to denote the order of *k*_0_-mers within *c* according to 𝒪_0_. Conditioned on any *c* with 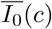, over the randomness of 𝒪_0_, the order of the *k*_0_-mers inside *c* now follows a random permutation of (2*w*_0_ + 2) = 2*w*, which we denote as ℛ(2).

Next, we consider fixing both *c* and 𝒪_0_ (the only randomness is in 𝒪), and calculate probability that the context is charged. Note that fixing *c* and 𝒪_0_ means fixed Ord(*c*), and fixed set of *k*-mers in 𝒞_0_. The order of the *k*-mers is still random due to randomness in 𝒪. For simplicity, if a *k*-mer is in 𝒞_0_, we call it a *UHS k-mer*. A *boundary UHS k-mer* is a UHS *k*-mer that is either the first or the last *k*-mer in the context *c*.

##### Lemma 14.

*Assume a fixed context c with no duplicate k*_0_*-mers and a fixed* 𝒪_0_. *Denote m*_*boundary*_ *as the number of boundary UHS k-mers (m*_*boundary*_ ∈ {0, 1, 2}*) and let m*_*total*_ *be the number of total UHS k-mers in the context. The probability that the context is charged, over the randomness of* 𝒪, *is m*_*boundary*_*/m*_*total*_.

*Proof.* We first note that *m*_total_ ≥ 1 for any context and any 𝒪_0_, due to 𝒞_0_ being a UHS over *k*-mers with path length *w*_0_ ≤ *w*, so the expression is always valid. Furthermore, there are also no duplicate *k*-mers as *k* > *k*_0_. As 𝒪 is random, every UHS *k*-mer in the window has equal probability to be the minimal *k*-mer. The context is charged if one of the boundary UHS *k*-mers is chosen in this process, and the probability is 1*/m*_total_ for each boundary UHS *k*-mer, so the total probability is *m*_boundary_*/m*_total_.

This proof holds for every *c* and 𝒪_0_ satisfying 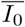. In this case, both *m*_boundary_ and *m*_total_ are only dependent on Ord(*c*). This means we can write the probability of charged context conditioned on 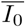 with a single source of randomness, as follows:

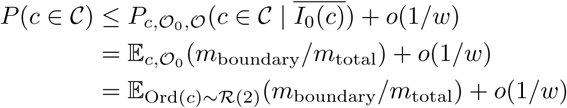

Next, we use *E*_0_ to denote the event that the first *k*-mer in the context is a UHS *k*-mer, and *E*_1_ to denote the event for the last *k*-mer. These two events are also only dependent on Ord(*c*). We then have *m*_boundary_ = **1**(*E*_0_) + **1**(*E*_1_). By linearity of expectation, we have the following:

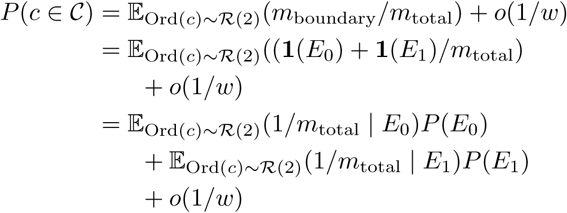

As the problem is symmetric, it suffices to solve one term. We have *P* (*E*_0_) = 2*/w*, because *E*_0_ is true if and only if the minimal *k*_0_-mer in the first *k*-mer is either the first or the last one, and there are *w k*_0_-mers in a *k*-mer. The only term left is 𝔼_Ord(*c*)∼ℛ(2)_(1*/m*_total_ |*E*_0_).

In the next two sections, we will upper bound this last term, which in turn bounds *P* (*c* ∈ 𝒞). It helps to understand why this argument achieves a bound better than a purely random minimizer, even though the Miniception looks very randomized. The context contains two UHS *k*-mers on average, because the relative size of 𝒞_0_ is 2*/w* + *o*(1*/w*), so it may appear the expectation term is close to 0.5, which leads to a density bound of 2*/w* + *o*(1*/w*), identical to a random minimizer. However, conditioned on *E*_0_, the context provably contains at least one other UHS *k*-mer, and with strictly positive chance contains two or more other UHS *k*-mers, which brings the expectation down strictly below 0.5.

#### 2.4.3 Deriving the Unconditional Distribution

In this section, we bound the quantity 𝔼_Ord(*c*)∼ℛ(2)_(1*/m*_total_ | *E*_0_) by deriving the distribution of *m*_total_, where Ord(*c*) is sampled from ℛ(2) conditioned on *E*_0_. We emphasize that at this point the actual *k*_0_-mers are not important and only their order matters. It is beneficial to view the sequence simply as the order Ord(*c*). To prepare for the actual bound, we will first derive the distribution of *m*_total_ assuming Ord(*c*) ∼ ℛ(2) without extra conditions.

We are interested in the asymptotic bound, meaning *w* → ∞, so we use the following notation. Let ℛ(*x*) denote the distribution of random order of *xw* elements. This is consistent with previous definition of ℛ(2), as a context contains 2*w k*_0_-mers. The *relative length* of a sequence is defined by its number of *k*_0_-mers divided by *w*. Given a sequence of relative length *x*, where the order of its constituent *k*_0_-mers follows ℛ(*x*), let *P*_*n*_(*x*) denote the probability that the sequence contains exactly *n* UHS *k*-mers. As a context is a sequence of relative length 2, we are interested in the value of *P*_*n*_(*x*) for *x* ≤ 2.

We derive a recurrence for *P*_*n*_(*x*). Fix *x*, the relative length of the sequence. We iterate over the location of the minimal *k*_0_-mer and let its location be *tw* where 0 ≤ *t* ≤ *x*.

There are two kinds of UHS *k*-mers for this sequence. The first kind contains the minimal *k*_0_-mer of the sequence, and there can be at most two of them: one starting with that *k*_0_-mer and one ending with that *k*_0_-mer. The second kind does not contain the minimal *k*_0_-mer, so it is either to the left of the minimal *k*_0_-mer or to the right of the minimal *k*_0_-mer, in the sense that it does not contain the minimal *k*_0_-mer in full. Precisely, it is from the subsequence that contains exactly the set of *k*_0_-mers left to the minimal *k*_0_-mer, or from the subsequence that contains exactly the set of *k*_0_-mers right to the minimal *k*_0_-mer: these two subsequences have an overlap of *k*_0_ − 2 bases but do not share any *k*_0_-mer, and neither contain the minimal *k*_0_-mer. We refer to these sequences as the left and right subsequence for conciseness.

This divides the problem of finding *n* UHS *k*-mers into two subproblems: finding UHS *k*-mers left of location *tw*, and finding UHS *k*-mers right of location *tw*. If we sample an order from ℛ(*x*), conditioned on the minimal *k*_0_-mer on location *tw*, the order of the *k*_0_-mers left of the minimal *k*_0_-mer follows ℛ(*t*), and similarly ℛ(*x* − *t*) for the *k*_0_-mers right of the minimal *k*_0_-mer. As we assume *w* → ∞, we ignore the fact that two subsequences combined have one less *k*_0_-mer. We prove in Supplementary Section S2.6 that this simplification introduces a negligible error. This means the subproblems have an identical structure to the original problem.

We start with the boundary conditions.

##### Lemma 15.

*For x* < 1, *P*_0_(*x*) = 1 *and P*_*n*_(*x*) = 0 *for n* ≥ 1. *For* 1 ≤ *x* ≤ 2, *P*_0_(*x*) = 2*/x* − 1.

*Proof.* A sequence with relative length less than 1 does not contain a single *k*-mer, so with probability 1 the sequence contains no UHS *k*-mer.

For a sequence with relative length between 1 and 2, define the middle region as the set of *k*_0_-mer locations that are at most *w* − 2 *k*_0_-mers away from both the first and the last *k*_0_-mer. The sequence contains no UHS *k*-mer if and only if the minimal *k*_0_-mer falls within the middle region, as only in this case every *k*-mer contains the minimal *k*_0_-mer but none have it at the boundary. The relative length of the middle region is 2 − *x*, as we assume *w* → ∞ (see Figure 3). As the order of *k*_0_-mers follows ℛ(*x*), every *k*_0_-mer has equal probability to be the minimal and it is in the middle region with probability (2 − *x*)*/x* = 2*/x* − 1.

**Figure 3:**
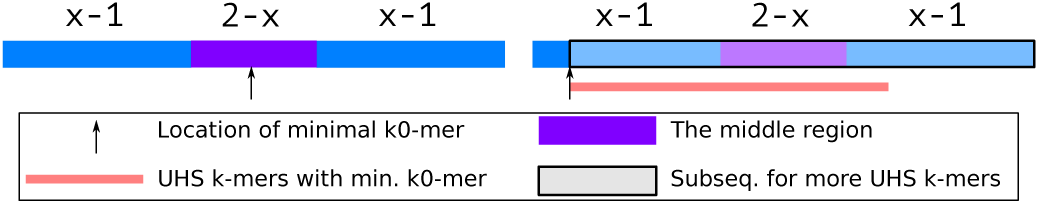
Setup for derivation of P_n_(x) with n ≥ 1 and 1 ≤ x ≤ 2. The text denotes the relative length of the corresponding subsequences. If the minimal k_0_-mer fall into the middle region (left panel), there are zero UHS k-mers in the sequence. Otherwise (right panel), there is at least one UHS k-mer with possibility for more from the subsequence.

For 1 ≤ *x* ≤ 2, we now derive the recurrence for *P*_*n*_(*x*) with *n* ≥ 1 (as seen in Figure 3). We define the middle region in an identical way as in last lemma, whose relative length is again 2 − *x*. If the minimal *k*_0_-mer is in the middle region, the sequence has exactly zero UHS *k*-mers. Otherwise, by symmetry we assume it is to the left of the middle region (that is, at least *w* − 1 *k*_0_-mers away from the last *k*_0_-mer in the sequence), with location *tw* where 0 ≤ *t* < *x* − 1. The sequence now always has one UHS *k*-mer, that is the *k*-mer starting with the minimal *k*_0_-mer, and all other (*n* − 1) UHS *k*-mers come from the subsequence right to the minimal *k*_0_-mer. The subsequence has relative length *x* − *t*, and as argued above, the probability of observing exactly (*n* − 1) UHS *k*-mers from the sub-sequence is *P*_*n*−1_(*x − t*). Averaging over *t*, we have the following:

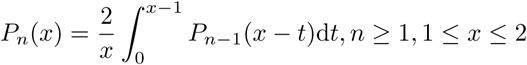

Given *P*_0_(*x*) = 2*/x* − 1, we can solve for the next few *P*_*n*_ for 1 ≤ *x* ≤ 2 as described in Supplementary Section S2.3. Recall our goal is to upper bound 𝔼_Ord(*c*)∼ℛ(2)_(1*/m*_total_ | *E*_0_). For this purpose, *P*_*n*_(*x*) is not sufficient as the expectation is conditioned on *E*_0_.

#### 2.4.4 Deriving the Conditional Distribution

We now define the events 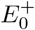 and 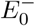. 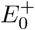 is the event that the first *k*-mer of the Miniception context is a UHS *k*-mer, because inside the first *k*-mer the minimal *k*_0_-mer is at the front. Similarly, 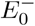 is the event where the first *k*-mer is a UHS *k*-mer, because the last *k*_0_-mer in the first *k*-mer is minimal. These events are mutually exclusive and have equal probability of 1*/w*, so 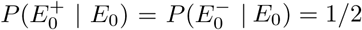.

##### Definition 8 (Restricted Distribution).

ℛ^+^(*x*) *for x* ≥ 1 *is the distribution of random permutations of xw elements, conditioned on the event that the first element is minimum among first w elements. Similarly*, ℛ^−^(*x*) *for x* ≥ 1 *is the distribution of random permutations of xw elements, conditioned on the event that the last element is minimum among first w elements.*

We now have the following:

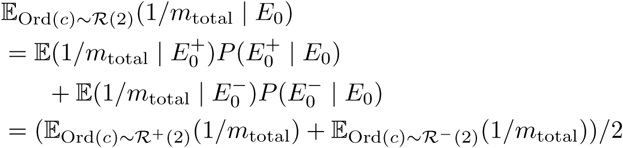

Based on this, we define 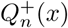 to be the probability that a sequence of relative length *xw*, where the order of *k*_0_-mers inside the sequence follows ℛ^+^(*x*), contains exactly *n* UHS *k*-mers. Our goal now is to determine 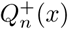 for *x* ≤ 2, which also bounds 𝔼_Ord(*c*)∼ℛ+(2)_(1*/m*_total_).

The general idea of divide-and-conquer stays the same in deriving a recurrence for 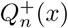. It is however trickier to apply this idea with a conditional distribution. We solve this issue by defining the following:

##### Definition 9 (Restricted Sampling).

*With fixed x and w, the restricted sampling process samples a permutation of length xw, then swap the minimum element in the first w element with the first element.*

##### Lemma 16.

*Denote the distribution generated by the restricted sampling process as* 𝒮^+^(*x*), *then* 𝒮^+^(*x*) = ℛ^+^(*x*).

We prove this in Supplementary Section S2.1. As the distributions are the same, we redefine 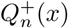 with 𝒮^+^(*x*). The boundary condition for *Q*^+^(*x*) is 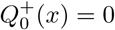 for all *x*, because the first *k*-mer is guaranteed to be a UHS *k*-mer (note that 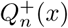 is defined only with *x* ≥ 1).

For *n* ≥ 1 and *x* ≤ 2, from the process of restricted sampling, we know with probability 1*/x* the minimal *k*_0_-mer in the sequence is the first *k*_0_-mer overall. In this case, the first *k*-mer is the only UHS *k*-mer that contains the minimal *k*_0_-mer, and all other UHS *k*-mers come from the subsequence without the first *k*_0_-mer whose relative length is still *x* as we assume *w* → ∞. We claim the following:

##### Lemma 17.

*Given an order of k*_0_*-mers sampled from* 𝒮^+^(*x*), *conditioned on the first k*_0_*-mer being overall minimal, the k*_0_*-mer order excluding the first k*_0_*-mer follows the unrestricted distribution* ℛ (*x*).

This is proved in Supplementary Section S2.1. This lemma gives the probability of observing (*n* − 1) UHS *k*-mers outside the first *k*_0_-mer of *P*_*n*−1_(*x*). Otherwise, we use the same argument as before by setting the location of the minimal *k*_0_-mer to be *tw*, where 1 ≤ *t* ≤ *x*. Only one UHS *k*-mer contains the minimal *k*_0_-mer with probability 1 (if *x* = 2, *t* = 2 happens with probability 0), and all other UHS *k*-mers come from the subsequence to the left of the minimal *k*_0_-mer. By a similar argument, the order of *k*_0_-mers within the left subsequence follows 𝒮^+^(*t*). These arguments are also shown in Figure 4. Averaging over *t*, we have the following recurrence for 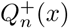, valid for 1 < *x* ≤ 2 and *n* ≥ 1:

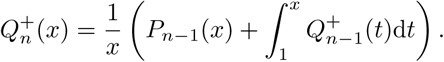

**Figure 4:**
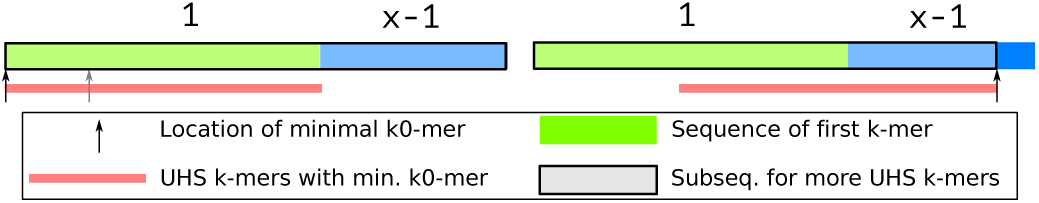
Setup for derivation of 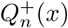 with n ≥ 1 and 1 ≤ x ≤ 2. The text denotes the relative length of the corresponding subsequences. If the minimal k_0_-mer is in the first k-mer, it will be the first k_0_-mer overall. There is one UHS k-mer with possibility for more in the subsequence without first k_0_-mer. Otherwise, the analysis is similar to the derivation of P_n_(x).

Replacing ℛ^+^(*x*) with ℛ^−^(*x*), we can similarly define and derive the recurrence for 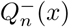 given 1 ≤ *x* ≤ 2. The process is highly symmetric to the previous case for 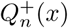, and we leave it to Supplementary Section S2.2. Similar to *P*_*n*_(*x*), we can derive the analytical solution to these integrals (see Section S2.3). By definition of 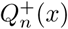, we now bound 𝔼_Ord(*c*)∼ℛ +(2)_ (1*/m*_total_) by truncating the distribution’s tail, as follows (omitting the condition for clarity):

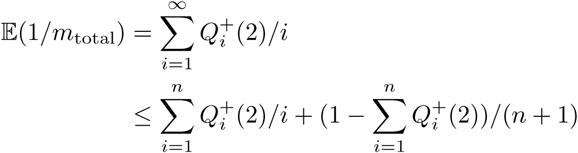

We can derive a similar formula for the symmetric term 𝔼_Ord(*c*)∼ℛ−(2)_(1/*m*_total_). For both *Q*^+^ and *Q*^−^, at *n* = 6 the tail probability 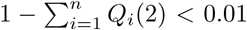, so we bound both terms using *n* = 6, resulting in the following:

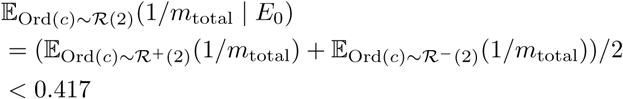

Finally, we bound the density of the Miniception, now also using the symmetry conditions (omitting the condition Ord(*c*) ∼ ℛ (2) for clarity):

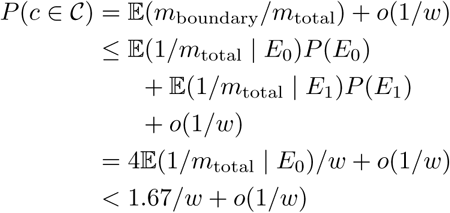

#### 2.4.5 Density Bounds beyond *x* = 2

We can derive the recurrence for 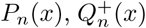 and 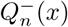 for *x* > 2, corresponding to the scenario where *w* ≈ (*x* − 1)*k* > *k*. By similar techniques, with suitably chosen *n*, we can upper bound the density of the Miniception from the values of 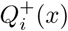 and 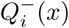 with *i* ≤ *n*. The resulting bound has form of *D*(*x*)*/w*+*o*(1*/w*), where *D*(*x*) is the density factor bound. The detailed derivations can be found in Section S2.4. We then have the following theorem:

##### Theorem 8.

*With x* ≥ 2, *w* = (*x*−1)(*w*_0_+1), *k* = *w*_0_+*k*_0_ *and k*_0_ > (3 + *ϵ*) log_*σ*_ (*x*(*w*_0_ + 1)), *the expected density of the Miniception is upper bounded by D*(*x*)*/w* + *o*(1*/w*).

## 3 Results

### 3.1 Asymptotic Performance of the Miniception

We use a dynamic programming formulation (Section S2.5) to calculate the density factor of the Miniception given *w* ≈ (*x* − 1)*k*, with a large value of *k* = 2500. As analyzed in Section S2.6, this accurately approximates *D*(*x*), which in turn approximates the density factor of the Miniception with other values of *k* up to an asymptotically negligible error. Figure 5 shows estimated *D*(*x*) for 2 ≤ *x* ≤ 8 with the described setup.

**Figure 5:**
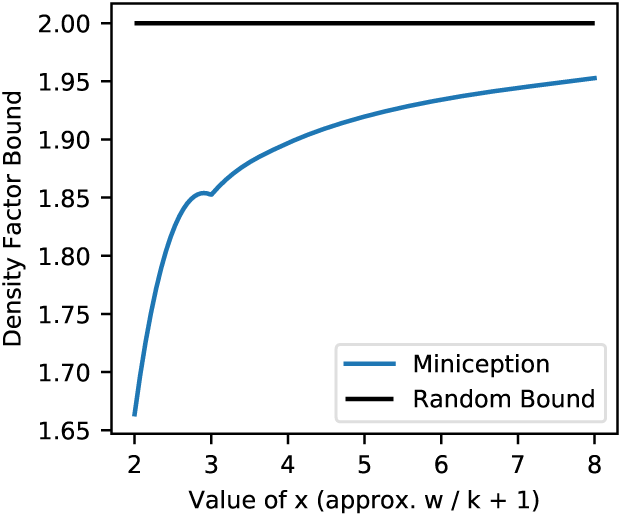
Density factor for the Miniception when w ≈ (x − 1)k and k = 2500. The random minimizer achieves density factor of constant 2 and is plotted for comparison.

Consistent with Section 2.4, *D*(2) ≈ 1.67, as *x* = 2 corresponds to the case *w* ≈ *k*. There is no analytical form for *D*(*x*) with *x* > 2, but this experiment suggests that as *x* grows, *D*(*x*) increases while staying below 2, the density factor of a random minimizer. That is, as *w* gets increasingly larger than *k*, the Miniception performance regresses to that of a random minimizer. We conjecture that *D*(*x*) = 2 − *o*(1) as *x* grows.

### 3.2 Designing Minimizers with Large *k*

As seen in the implementation of Miniception (Section S3), the run time of the Miniception minimizer is the same as a random minimizer. Therefore, it can be used even for large values of *k* and *w*. This contrasts to PASHA (Ekim *et al.* (2020)), the most efficient minimizer design algorithm, which only scales up to *k* = 16.

We implemented the Miniception and calculated its density by sampling 1 000 000 random contexts to estimate the probability of a charged context (which is equivalent to estimating the density as shown in Lemma 2). We show the results for Miniception against lexicographic and random minimizers for 10 ≤ *k* ≤ 30, and *w* = {10, 100} (Figure 6). These parameter ranges encompass values used by bioinformatics software packages. The Miniception has a single tunable parameter *k*_0_. We tune this parameter in our experiments by either setting *k*_0_ = *k* − *w*, or choosing the best performing *k*_0_ from 3 ≤ *k*_0_ ≤ 7.

**Figure 6:**
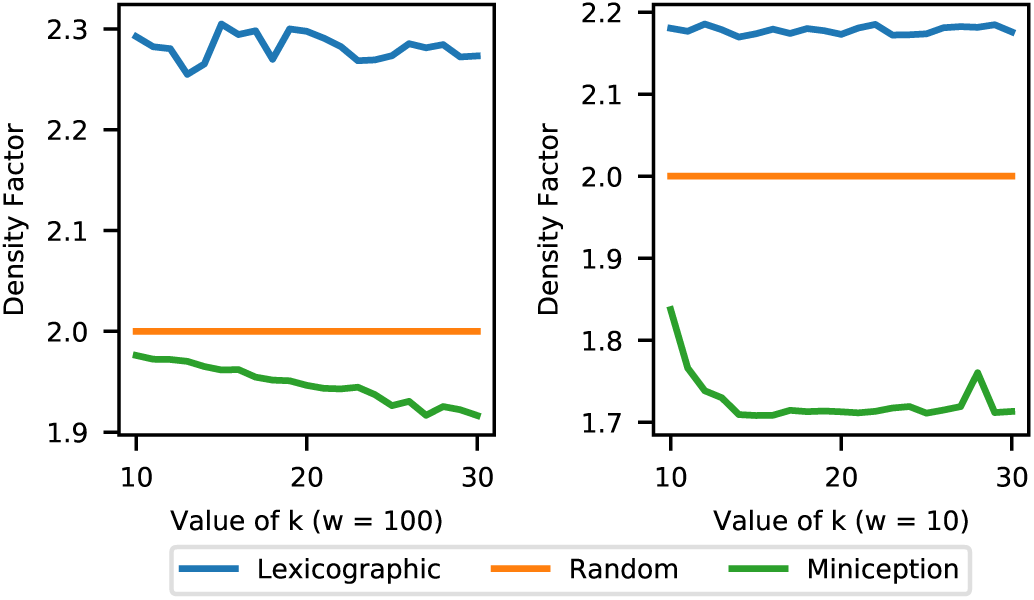
Comparing density of the Miniception against lexicographic and random minimizers. We experiment with w = 10 (left half) and w = 100 (right half).

The Miniception consistently performs better than the lexicographic and random minimizers in all tested scenarios. For *w* = 10, *k* is larger than *w* and the Miniception achieves density factor of ≈ 1.72 for *k* ≥ 13. Given that 1*/w* = 0.1, our bound on the density factor of 1.67 + *o*(1*/w*) holds relatively well for these experiments.

For *w* = 100, *w* is larger than *k* and we observe the same behavior as in Section 3.1: the performance degrades when *k* becomes smaller than *w*. Our theory also correctly predicts this behavior, as the decrease of *x* ≈ *w/k* + 1 improves the density bound as seen in Section 2.4.

### 3.3 Comparison with PASHA

In Figure 7, we compare the Miniception with PASHA. We downloaded the PASHA generated universal hitting sets from the project website, and implemented the compatible minimizers according to Ekim *et al.* (2020). Our test consists of two parts. For the first part, we fix *k* = 13 and vary *w* from 20 to 200. For the second part, we fix *w* = 100 and vary *k* from 7 to 15. This setup features some of the largest minimizers designed by PASHA, the state-of-the-art in large minimizer design (we are unable to parse the UHS file for *k* = 16).

**Figure 7:**
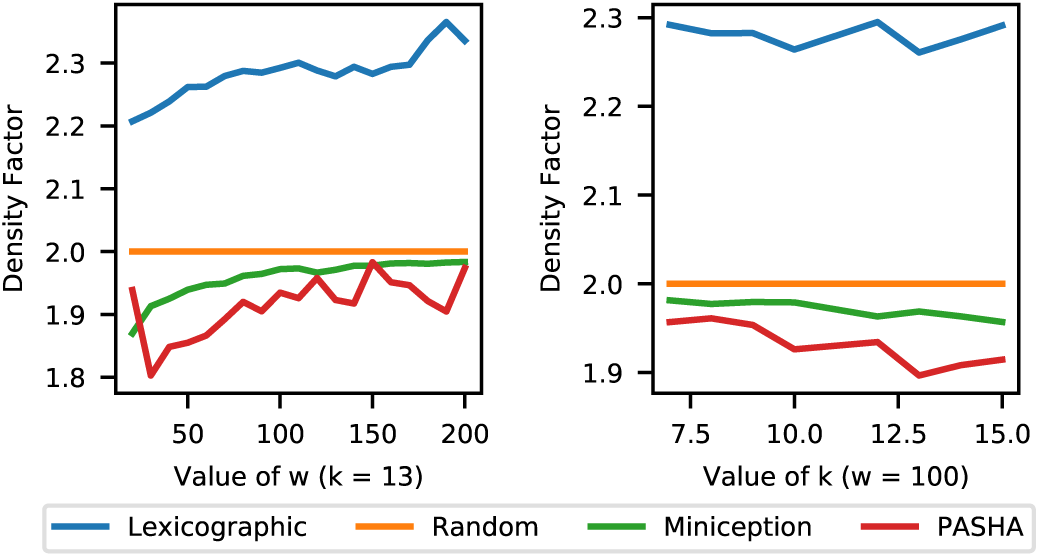
Density factor comparison, now also with minimizers derived from PASHA outputs. Left half: experiments with fixed k = 13 and varying w. Right half: experiments with fixed w = 100 and varying k.

The Miniception still performs better than the random minimizers for these configurations, but PASHA, even though it is a heuristic without a density guarantee, overall holds the edge. We also perform experiments on the hg38 human reference genome sequence, where we observe similar results with a smaller performance edge for PASHA, as shown in Supplementary Section S4.

## 4 Discussion

### 4.1 Limitations and Alternatives to Minimizers

The 1*/w* bound is not the only density lower bound for minimizers. Specifically, Marçais *et al.* (2018) proved the following lower bound:

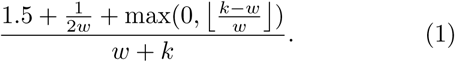

As *w* grows compared to *k*, this implies that the density factor of the minimizers are lower bounded by a constant up to 1.5. We also observed that performance of minimizers, both the Miniception and the PASHA compatible ones, regress to that of a random minimizer when *w* increases. Unfortunately, this is inherent to minimizers. With a fixed *k*, as the window size *w* grows, the *k*-mers become increasingly decoupled from each other and the ordering 𝒪 plays less of a role in determining the density.

The minimizers are not the only class of methods to sample *k*-mers from sequences. Local schemes are generalizations of minimizers that are defined by a function *f* : Σ^*w*+*k*−1^ → {0, 1, …, *w* − 1} with no additional constraints. They may not be limited by existing lower bounds ((1) for example), and developing local schemes can lead to better sampling methods with densities closer to 1*/w*.

### 4.2 Perfect Schemes and Beyond

This work answers positively the long standing question on the existence of minimizers that are within a constant factor of optimal. Even though the original papers introducing the winnowing and minimizer methods proposed a density of 2*/w*, their analysis only applied to particular choices of *k* and *w*. Theorem 2 and 3 give the necessary and sufficient conditions for *k* and *w* to be able to achieve *O*(1*/w*) density. As a direct consequence, Theorem 2 gives the first constant factor approximation algorithm for a minimum size UHS, which improves on our previous result of a ln(*w*) factor approximation (Zheng *et al.*, 2020). These theorems also settle the question on existence of asymptotically perfect forward and local schemes.

In general, studies on the asymptotic behavior of minimizers have proven very fruitful to deepen our understanding of the minimizers and of the associated concepts (structure of the de Bruijn graph, decycling sets and universal hitting sets). However, there is a sizable gap between the theory and the practice of minimizers.

One example of this gap is the way we prove Theorem 2: the density of the lexicographic minimizers reaches *O*(1*/w*) whenever it is possible for any minimizer. This means the lexicographic minimizers are optimal for asymptotic density. However, in practice, they are usually considered the worst minimizers and are avoided. Another example is the fact that heuristics such as PASHA, while unable to scale as our proposed methods and being computationally extensive, achieves better density in practice (for the set of parameters it is able to run on) with worse theoretical guarantee. Now that we have mostly settled the problem of asymptotical optimality for minimizers, working on bridging the theory and the practice of minimizers is an exciting future direction.

The core metric for minimizers, the density, is measured over assumed randomness of the sequence. In many applications, especially in bioinformatics, the sequence is usually not completely random. For example, when working with a read aligner, the minimizers are usually computed on a reference genome, which is known to contain various biases. Moreover, this sequence may be fixed (e.g., the human genome). In these cases, a minimizer with low density in average may not be the best choice. Instead, a minimizer which selects a sparse set of *k*-mers specifically on these sequences would be preferred. The idea of “sequence specific minimizers” is not new (e.g., see DeBlasio *et al.* (2019); Chikhi *et al.* (2015b)), however, it is still largely unexplored.

## S1 Asymptotically Optimal Lexicographical Minimizers

In this section, we prove that the asymptotical optimality for the lexicographic minimizer with *k*_0_ = ⌊log_*σ*_ (*w/*2)⌋ − 2 can be used to derive the asymptotical optimality for other values of *k* ≥ log_*σ*_(*w*) − *c*.

### Lemma 8.

*The lexicographic minimizer with k* = *k*_0_ − *c for constant c* ≥ 0 *has density O*(1*/w*).

*Proof.* We define *W*′^+^ and *W*′^−^ in the same way for this new minimizer with parameter *k*. We also call the new minimizer as *k*-minimizer, and the minimizer in Theorem 1 as *k*_0_-minimizer.

We now claim |*W*′^+^| ≤ |*W*^+^|. This is because given a window in *W*′^+^, we can always append 0^*c*^ before the start of the window to get a window (for the *k*_0_-minimizer) in *W*^+^. Assume otherwise, we let *z*_0_ = 0^*c*^ … be the *k*_0_-mer at the start of the extended window and *z*_1_ be the *k*_0_-mer picked by the *k*_0_-minimizer. This requires *z*_1_ to also have form of 0^*c*^ …, and that the *k*-mer after 0^*c*^ is strictly smaller than the last *k*-mer of *z*_0_. However, if this is the case, the original window will not be in *W*′^+^ as the first *k*-mer is not minimal.

Similarly, we claim |*W*′^−^| ≤ |*W*^−^| as given a window in *W*′^−^ we can append 0^*c*^ after the end of the window to get a window in *W*^−^. Assume otherwise, we let *z*_0_ = … 0^*c*^ be the last *k*_0_-mer at the end of the extended window and *z*_1_be the *k*_0_-mer picked by the *k*_0_-minimizer. Since *z*_0_ ends with 0^*c*^ and *z*_1_ is smaller or equal to *z*_0_, the *k*-prefix of *z*_1_ must be smaller or equal to the *k*-prefix of *z*_0_. This means the original window will not be in *W*′^−^ as there is a *k*-mer that is smaller or equal to the last *k*-mer.

Therefore |*W*′^+^| = *O*(*σ*^*w*^) and |*W*′^−^| = *O*(*σ*^*w*^), and the density is *σ*(|*W*′^+^| + |*W*′^−^|)*/σ*^*w*+*k*^ = *O*(*σ*^−*k*^) = *O*(1*/w*).

On the other hand, for large *k* the following lemma establishes asymptotically optimal density. Recall *W*^*^ is the set of windows (for the *k*_0_-minimizer) such that the last *k*_0_-mer is among the smallest, but not necessarily unique (so the minimizer might not always pick it).

### Lemma 9.

*The lexicographic minimizer with k* > *k*_0_ *has density O*(1*/w*).

*Proof.* We define *W*′^+^ and *W*′^−^ in the same way for this new minimizer, and again call the new minimizer the *k*-minimizer.

We first claim 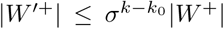. This is because given a window in *W*′^+^, we can remove the last *k* − *k*_0_ bases to get a window in *W*^+^. To see this, we only need to show for the shorter window, the first *k*_0_-mer is less than or equal to every other *k*_0_-mer. Since the original window is in *W*′^+^, the first *k*-mer in the original window is less than or equal to every other *k*-mer. This means the *k*_0_-long prefix of the first *k*-mer is also less or equal to the *k*_0_-long prefix of every other *k*-mer in the original window, which is exactly the condition for the short window in *W*^+^. For every window *x* in *W*^+^, there are 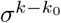 windows in Σ^*w*+*k*−1^ that *x* is a prefix of, which means 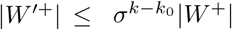.

We now claim 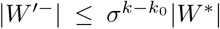. Similarly, given a window in *W*′^−^, we can again remove the last *k* − *k*_0_ bases to get a window in *W*^*^. As the original window is in *W*′^−^, the last *k*-mer in the original window is less than every other *k*-mer. This means the *k*_0_-long prefix of the last *k*-mer is less or equal to the *k*_0_-long prefix of every other *k*-mer in the original window. (Note that if a *k*-mer is less than another, their prefix can be equal.) The *k*_0_-long prefixes of *k*-mers in the original window are exactly the *k*_0_-mers of the shorter window, so the short window is in *W*^*^. With a similar argument, we have 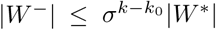.

These two facts combined means we can calculate the density of the new minimizer as follows:

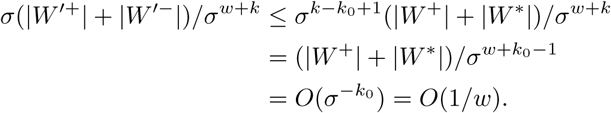

## S2 More Analyses of the Miniception

This section contains analyses of the Miniception that are not included in the main text.

### S2.1 Restricted Sampling

In this section, we formally prove the correctness of the restricted sampling process.

#### Lemma 16.

*Proof.* We prove the two distributions are identical in two parts. First, we show they have the same support (generate the same set of permutations). Note that any permutations sampled from 𝒮^+^(*x*) satisfies 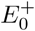 by the swapping process, and any permutation satisfying 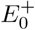 can be sampled from 𝒮^+^(*x*) as it can simply be the initial distribution and unchanged by the swapping process. Second, given ℛ^+^(*x*) is a uniform distribution over permutations satisfying 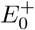, we only need to prove for any two permutation satisfying 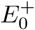 they are generated by 𝒮^+^(*x*) with the same probability. This is true because every permutation satisfying 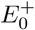 has exactly *w* preimages in the swapping process.

#### Lemma 17.

*Given an order of k*_0_*-mers sampled from* 𝒮^+^(*x*), *conditioned on the first k*_0_*-mer being overall minimal, the k*_0_*-mer order excluding the first k*_0_*-mer follows the unrestricted distribution* ℛ(*x*).

*Proof.* Our goal is to prove the two distributions are equal, and we will use the same two-step process. First, every permutation can be generated by both processes. Second, as proved before, 𝒮^+^(*x*) generates each eligible permutation with identical probability, and this holds for the conditional distribution as each permutation has exactly one preimage. This means it is the same distribution as ℛ(*x*) where each permutation is also generated with the same probability.

### S2.2 Analysis of 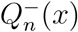 with *x* ≤ 2

Based on our analysis of 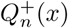 for *x* ≤ 2, as presented in the main text, we derive the recurrence for 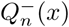 for *x* ≤ 2 here.

To start with, we similarly define the restricted sampling process (where we swap the minimal element in the first *w* elements with the *w*^th^ element instead). We define 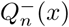 to denote the same quantity as 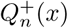, except the order is now sampled from the new restricted sampling process 𝒮^−^(*x*). The derivation of 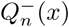 is identical when the minimal *k*_0_-mer in the sequence is not the *w*^th^ one. With probability 1*/x*, the minimal *k*_0_-mer will be the *w*^th^ one. In this case, if *x* < 2, again only one UHS *k*-mers will contain the minimal *k*_0_-mer, but with *x* = 2 two *k*-mers will contain it (with *x* = 2, the *k*_0_-mer is in the middle of the sequence, with at least *w* − 1 *k*_0_-mers away from each end). All other UHS *k* _0_-mers now come from the subsequence right to the minimal *k*_0_-mer, whose relative length is now *x* − 1, and similar to our previous argument, the order within the subsequence follows ℛ(*x* − 1). This yields the following recurrence:

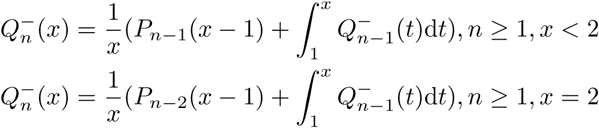

### S2.3 Analytical Solution for 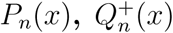 and 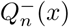 at *x* ≤ 2

In this section, we provide the analytical solution to the integrals given *x* ≤ 2, up to *n* = 6 as follows. The formula of *P*_*n*_(*x*) for 4 ≤ *n* ≤ 6 is more complicated and is omitted here for clarity. We provide (in our Github repository) automatic symbolic integration codes that computes these integrals up to any specified *n*, and calculate *D*(2) based on the integrals.

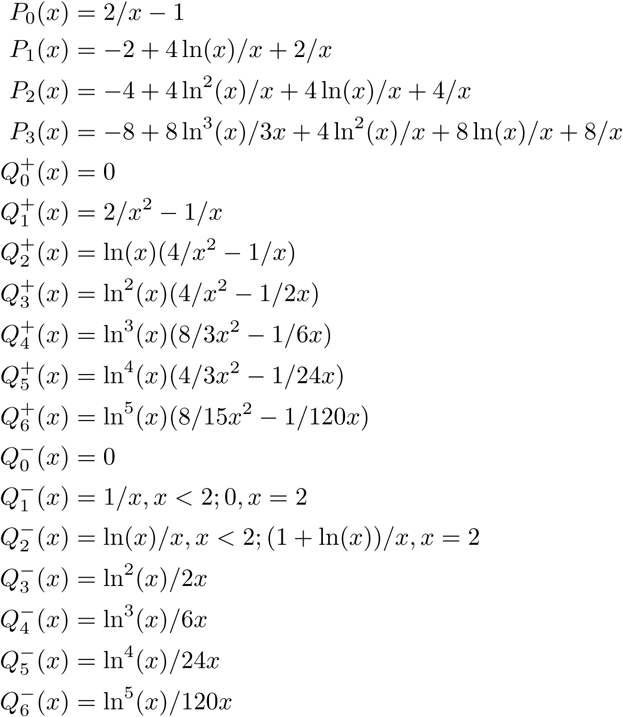

### S2.4 Deriving the Integrals for *x* > 2

To extend our analysis to other values of *w*, we will derive the recurrence formula for 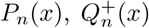 and 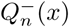 for *n* ≥ 2. Note that as *w* = *w*_0_ + 1 no longer holds, the relative length of a sequence is defined as the number of *k*_0_-mers in the sequence divided by *w*_0_+1, and ℛ(*x*) is now a random permutation of *x*(*w*_0_ + 1) elements. Definition for 𝒮^±^(*x*) and the desired quantities change accordingly.

We will follow the same general argument as before, by iterating over the location of minimal *k*_0_-mer within the sequence as *t*(*w*_0_ + 1) for 0 ≤ *t* ≤ *x*, and discuss the different scenarios. The extra consideration comes from the fact that now it is possible both left and right subsequences can produce UHS *k*-mers, so we also need to iterate over the number of UHS *k*-mers from one subsequence. We will define the convolution operator to represent this iteration process:

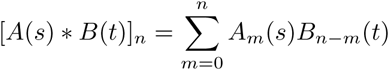

We start with the derivation of *P*_*n*_(*x*) for *x* > 2. For *n* = 0 and *x* > 2, *P*_*n*_(*x*) = 0 as wherever the minimal *k*_0_-mer is, there will be one UHS *k*-mer containing it. For the rest, we define the middle region as the subregion that is at least *w*_0_ *k*_0_-mers away from both the first and the last *k*_0_-mer in the sequence. Its relative length is *x* − 2. If the minimal *k*_0_-mer falls in this region, two UHS *k*-mers (recall a *k*-mer consists of (*w*_0_ + 1) *k*_0_-mers) will contain the *k*_0_-mer and both subsequences split by the minimal *k*_0_-mer may contain more UHS *k*-mers. If the minimal *k*_0_-mer falls outside this region, the analysis is identical to the previous case. This yields the following recurrence:

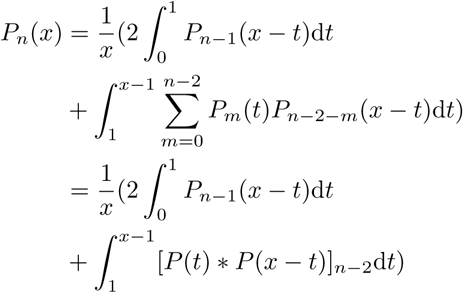

For 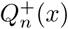, we similarly define the middle region. If the minimal *k*_0_-mer before swapping process falls to the left of the middle region (with probability 1*/x*), it falls within the first *k*-mer and its order is swapped with the first *k*_0_-mer. The probability of observing *n* − 1 UHS *k*-mers outside the first *k*_0_-mer is the same *P*_*n*−1_(*x*). If the minimal *k*_0_-mer is to the right of the middle region, one UHS *k*-mer contains this *k*_0_-mer and all other UHS *k*-mers comes from the left subsequence, following 𝒮^+^(*x* − 1). If the minimal *k*_0_-mer is in the middle region, two UHS *k*-mers contain the *k*_0_-mer and both subsequences may contain more UHS *k*-mers, with the order in the left subsequence conditioned on 𝒮^+^(*t*) and the order in the right subsequence conditioned on ℛ(*x* − *t*). This yields the following recurrence:

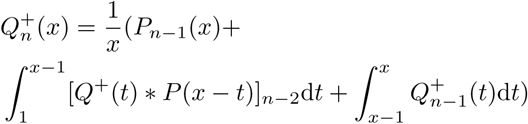

The derivation for 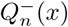 is extremely similar to that of 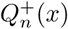. Note that now *x* ≥ 2, two UHS *k*-mers are guaranteed when the minimal *k*_0_-mer is the *w*^th^ one.

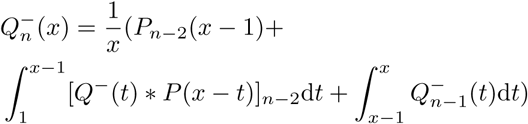

Following the methods of truncating distributions outlined in the main text, noting that now 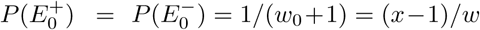, we can calculate density factor bound as follows:

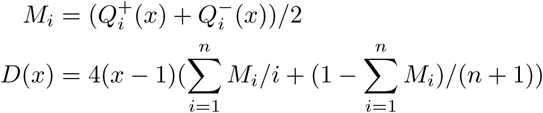

assuming the integrals are derived up to *n* UHS *k*-mers. The final density bound will be *D*(*x*)*/w* + *o*(1*/w*).

### S2.5 Estimating *D*(*x*) with Dynamic Programming

As *D*(*x*) represents the density of the Miniception when *w* ≈ (*x* −1)*k*, with *k* → ∞, it is natural to approximate *D*(*x*) by selecting a large value of *k* and calculate the density of the Miniception with *w* = (*x* − 1)*k*. Here we make the assumption that all *k*_0_-mers in the string are unique and *k*_0_ ≪ *k*, so we simply take *k*_0_ = 1, *w* = (*x* − 1)*k* and assume the order of the *k*_0_-mers in the context follows ℛ(*x*). By our analysis for the integral method (See Section S2.6), the final density bound will be off by *O*(1*/k*) compared to the integrals, so by choosing *k* to be large enough we can ensure the derived bound is close to the integral bound, which again by Section S2.6 approximates the behavior of the Miniception for arbitrary large values of *k*.

Now, let *P*[*l, n*] be the probability that given permutation of *l k*_0_-mers, there are exactly *n* UHS *k*-mers. Recall a *k*-mer is a UHS *k*-mer if its first or last *k*_0_-mer is the smallest in the subsequence, which is of length *k* now. We again enumerate over the location of the smallest *k*_0_-mer in permutation, and denote this value *i*. Each specific location is the minimum with probability 1*/l*. If *i* ≥ *k* − 1, the *k*-mer that ends at *i* (meaning its last *k*_0_-mer is this one) is a UHS *k*-mer. By symmetry, if *i* ≤ *l* − *k*, the *k*-mer starting at *i* is a UHS *k*-mer. The rest of UHS *k*-mers will come from the two subsequences, obtained by removing the *i*^th^ *k*_0_-mer from the sequence, and as before, we need to enumerate the number of UHS *k*-mers from either sub-sequences. This results in the following recurrence, where we use *f*_*l,n*_(*i*) to denote the number of UHS *k*-mers that need to come from either subsequences:

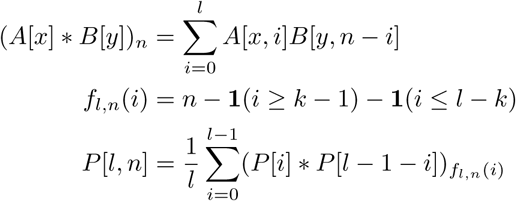

With the dynamic programming, we only need to provide that *P*[*i*, 0] = 1, *P*[*i, n*] = 0 for *i* < *k* and *n* ≥ 1. We can then calculate the values of *P*[*l, n*] up to *l* = *xk* and some preset value of *n* with the requirement *n* > 2*x*.

We can similarly generate the recurrence relationship for *Q*^+^[*l, n*] and *Q*^−^[*l, n*] (derivations are highly similar and omitted here):

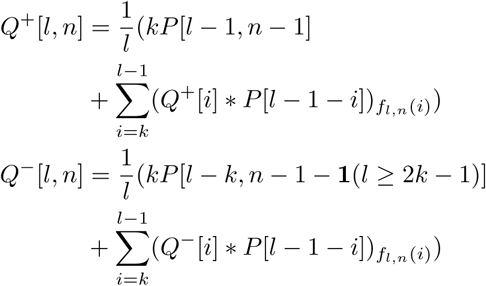

The approximate density factor bound (we use the density factor bound for numerical stability and ease of comparison), which we denote as *D*_*k*_(*x*), can be calculated as follows using our previous argument of truncating the distribution.

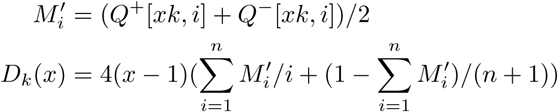

The final density bound for the Miniception under this configuration will be *D*_*k*_(*x*)*/w* +*o*(1*/w*), where the *o*(1*/w*) term comes from the event that some *k*_0_-mers in a context can be identical.

### S2.6 Error Analysis for the Integral Method

In the derivation for the Miniception, we assume *k* → ∞ and use an integral instead of summation for derivation of *P*_*n*_(*x*) and other quantities. In this section, we show the error introduced by this method is small enough to not affect the final result asymptotically. Specifically, we show the following:

#### Lemma S18.

*Let D*(*x*) *be the density factor bound derived from the integrals, and D*_*k*_(*x*) *be the density factor bound calculated from the dynamic programming with window length* (*x*−1)*k and k-mer length k (measured in number of k*_0_*-mers), assuming all k*_0_*-mers are distinct. We have* |*D*(*x*) − *D*_*k*_(*x*)| = *O*(*nx/k*), *where n is the largest number of UHS k-mers considered by both processes.*

As shown in the main text, this lemma serves a dual purpose. It means *D*(*x*) is a good approximation to any *D*_*k*_(*x*) up to asymptotically negligible errors, and by calculating *D*_*k*_(*x*) for sufficiently large *k*, we can estimate *D*(*x*) accurately which in turn approximates other *D*_*k*_(*x*). In other words, *D*(*x*) serves as a bridge to connect the density factor bound *D*_*k*_(*x*) for different values of *k*, which then bound the density of the actual minimizer as long as *k*_0_ satisfies the condition for Lemma 8. Note that in derivation of *D*_*k*_(*x*) the concept of *k*_0_-mers are already abstracted away, and for simplicity we assume *k*_0_ = 1 in that process, meaning every *k*-mer now consists of *k k*_0_-mers.

Recall in the main text we set up the integrals to derive the distribution of *m*_total_ conditioned on first *k*-mer being a UHS *k*-mer. Following the convention in the dynamic programming process, we assume *k*_0_ = 1, meaning *k* = *w*_0_ + 1. The context has exactly *xk k*_0_-mers, and each *k*-mer is of length *k*. Note that all lengths in the section are measured in *k*_0_-mers. As shown in the main text, conditioned on *E*_0_, the context will contain at least two UHS *k*-mers.

Recall that we derive the values of *P*_*n*_(*x*) up to *n*. An alternative view of the integral recurrence is that we determine at most *n* + 1 locations that contains the minimal *k*_0_-mers in a certain subsequence, and calculate the number of UHS *k*-mers based on the order and the distance of these locations. For example with *x* = 2, if the first location (the minimal *k*_0_-mer in the whole context) is on the left half of the context, the second location is for the minimal *k*_0_-mer in the right subsequence. If the second location is in the middle region of that subsequence, we determined the number of UHS *k*-mers (1 in this case) using two locations. If instead, the second location is not inside the middle region, a third location is determined inside the sub-sub-sequence that could still generate a UHS *k*-mer, and these three locations can collectively determine how many UHS *k*-mers are in the whole context. This process continues until no subsequences can generate more UHS *k*-mers, or *n* + 1 UHS *k*-mers has been generated. There are only *n* + 1 determined locations total, as for each recursion there is one corresponding UHS *k*-mer, either right before the start of the subsequence or right after the end of the subsequence. This is similar when we derive the values of 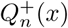 and 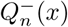, where we determine the location of the minimal *k*_0_-mer before swapping.

We denote the sequential decision process to determine number of UHS *k*-mers, conditioned on 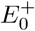 (corresponding to 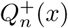) and 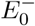 (corresponding to 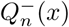), as 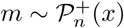 and 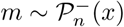, respectively. Formally:

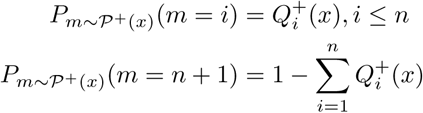

And this is symmetric for 𝒫^−^(*x*). With this definition, we have:

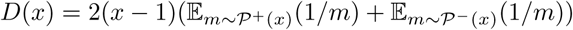

Similarly, we can propose an alternative view of the dynamic programming, where we determine at most *n* + 1 locations that contains the minimal *k*_0_-mers in a certain sequence. The difference is that in the previous cases, the locations are picked on a real axis [0, *x*], but now it is picked from a discrete set of locations in [*xk*]. We will now argue the two processes are not that different.

We first establish a mapping *M*(*t*) : ℝ → [*xk*] from real numbers to discrete locations, by multiplying the real number by *k* and rounding down. When a real number is sampled uniformly from [0, *x*], its mapping will be a uniform distribution over [*xk*] (that is, 1*/xk* chance to map into each location). Now, we consider emulating the decision process from the dynamic programming (which we denote 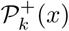), for simplicity). Each time a location is randomly sampled (the location of minimal *k*_0_-mer in current subsequence of consideration) from 𝒫^+^(*x*), we emulate one step of 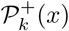 by selecting the mapped discrete location as the location of the minimal *k*_0_-mer. The process continues until the end of decision process.

We now examine the emulation process in more detail. At each step of the decision process, we sample the location of the minimal *k*_0_-mer in current subsequence of consideration, before the swapping process if the subsequence of consideration is still conditioned on 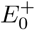 or 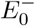. The boundary of the subsequence is determined by two previous samples (or simply the boundary), which we denote *L* and *R*. The discrete decision process we are emulating will be sampling the location of the minimal *k*_0_-mer in current subsequence of consideration, which will be the subsequence from *M*(*L*) to *M*(*R*), including neither ends. It can be seen that we are sampling from a slightly longer range in the original process. As long as our sampled location *T* satisfies *M*(*L*) < *M*(*T*) < *M*(*R*), the distribution of *M*(*T*) is uniform over possible selections. If *M*(*T*) = *M*(*L*) or *M*(*T*) = *M*(*R*) at any step of the emulation, we call the whole process failed and do not continue. The swapping process can be addressed similarly, as in this case *L* = 0 is guaranteed, and the emulated and original decision process will never disagree on whether the swap should be triggered.

Assume *M*(*T*) passes this check, meaning it is uniformly sampled from possible locations between *M*(*L*) and *M*(*R*). We will now decide on the next step in the process. This includes two parts: Counting number of UHS *k*-mers including the minimal *k*_0_-mer, and determining if a subsequence can still produce more UHS *k*-mers. Both parts involves two distance checks: If *T* − *L* > 1 in the original decision process, and if *M*(*T*) − *M*(*L*) ≥ *w*_0_ in the discrete decision process, for checking if the *k*-mer ending with the minimal *k*_0_-mer in this subsequence constitutes a valid UHS *k*-mer. If the original decision process and the emulated decision process will give different answer to this check, we call the whole process failed and do not continue. For deciding if the left subsequence can produce more UHS *k*-mers, we replace *w*_0_ with *w*_0_ + 1 and the analysis is the same. It is also symmetric for the right subsequence and checking the *k*-mer starting with minimal *k*_0_-mer.

Now, if a decision process does not fail (by either of the previously defined criteria), we say it perfectly emulates a discrete decision process. By the way it is defined, the emulated process randomizes the location of minimal *k*_0_-mers correctly, make the same decision as the original process at every step, and count the same number of UHS *k*-mers. We now characterize these “safe decision processes” with the following notation. Let 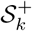 denote a distribution whose support is, {×, 1, 2, 3, …, *n* + 1} derived from the aforementioned emulation process. × denotes a failed process, and an integer denotes the number of UHS *k*-mers determined from the process. We claim 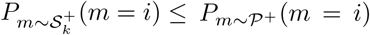 and 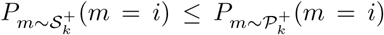 simultaneously for all *i* ∈ {1, 2,, 3, …, *n* + 1}. Intuitively, the distribution 𝒫^+^ and 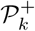 looks the same if the emulation is successful.

The first part of the statement can be proved by the definition of the emulation process: 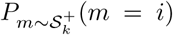 is the probability that the emulation successes and returns *i* UHS *k*-mers, which is never greater than the probability of *i* UHS *k*-mers regardless of the emulation outcome (*P* (*A*∩*B*) ≤ *P* (*A*)). The second part is can be proved by viewing the emulation from the perspective of the discrete process (which we call a reverse emulation, however they are in fact the same process): At each step of the discrete decision process, we first randomly select the location of the minimal *k*_0_-mer, and randomly choose a real number that maps to this location as its real coordinate. We then do a failure check by rolling a random number inside [*L, R*] (from the real number locations selected in previous steps), and fail the process if the rolled location collide. Similarly, at each distance check we first get the outcome from discrete locations, then fail the process if the real number locations disagree with the outcome. In this perspective, we can prove the second part of the statement in the same way: 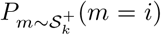 is the probability that this reverse emulation successes and returns *i* UHS *k*-mers, and is never greater than the probability of getting *i* UHS *k*-mers regardless of the outcome of reverse emulation.

We now bound the probability of failure, that is 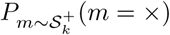. As the decision process consists of *n* + 1 steps, we will bound the probability at each step. At each step, the interval for random selection has at least relative length of 1 (otherwise no UHS *k*-mers will be from this section), and there are at least *k k*_0_-mers for selection. The key observation here is there are only 6 ways to fail (colliding with *M*(*L*), disagreement on whether left subsequence contains more UHS *k*-mers, disagreement on whether the *k*-mer ending at smallest *k*_0_-mer is valid UHS *k*_0_-mer, and their symmetric counterparts), and for each of them, there is a interval of length at most 1*/k* such that the fail event is triggered if and only if the minimal *k*_0_-mer is inside the interval. In other words, the probability to fail at each decision step is at most 6*/k* = *O*(1*/k*), and the probability of failure at any step is *O*(*n/k*).

We now have the tools to bound |*D*(*x*) − *D*_*k*_(*x*)|. We first bound the term 𝔼_*m*∼𝒫+(*x*)_(1/*m*). We denote *S* as the event that the emulation is successful, and ℱ^+^(*x*) as the distribution of UHS *k*-mer count when the emulation fails.

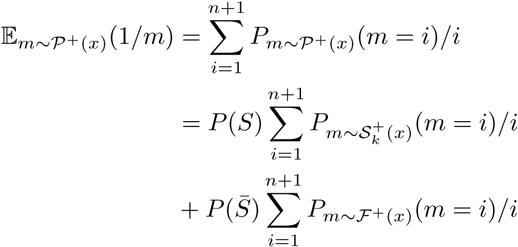

And similarly, let 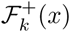 as the distribution of UHS *k*-mer count when the reverse emulation fails.

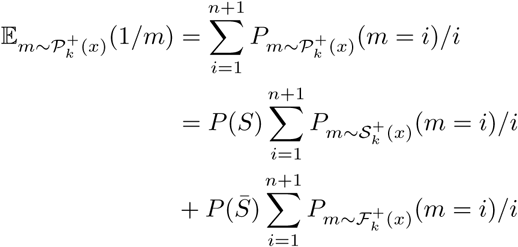

This leads to the following:

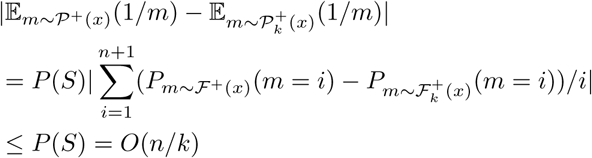

Substitute this back, we get |*D*(*x*) − *D*_*k*_(*x*)| = *O*(*nx/k*).

Note that this bound is not tight. This is because we overestimate the difference term 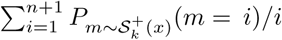 by simply upper bounding it by 1. However, for larger value of *k* and *x*, the context is almost guaranteed to have around 2*x* UHS *k*-mers, and the difference here is more on the order of *O*(1*/x*^2^). This means in practice the bound is more like *O*(1*/k*) (considering *n* = Θ(*x*) is a reasonable choice) with the possibility of even smaller (as we do not consider that positive terms and negative terms cancel out each other), which also explains why we can pick *k* = 2500 in our simulations.

## S3 Implementing the Miniception

The implementation takes a sequence *S* as input (as well as other parameters specifying the minimizer), and returns the list of picked locations. We will derive a linear time implementation, meaning the algorithm runs in time *O*(*k* |*S*|) as it takes *O*(*k*) time just to process and compare *k*-mers. We will discuss our algorithms based on the assumption that a *k*-mer fits in a word (This means *k* ≤ 32 for 64-bit systems), Value of *k* beyond this limit is rare in practice, and our algorithm also easily adapts to general values of *k*.

While the Miniception is defined with many randomness, implementing it means sticking to one “instantiation”, and the orders 𝒪_0_ and 𝒪 are fixed. We will assume the alphabet is {0, 1, 2, 3} so a *k*-mer can be written as an integer in [4^*k*^]. We assume constant time access to 𝒪_0_ as a function 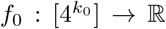, such that *x*_0_ < *x*_1_ in 𝒪_0_ if and only if *f*_0_(*x*_0_) < *f*_0_(*x*_1_), and similarly a function *f* : [4^*k*^] → ℝ for some order function that agrees with 𝒪 when restricted to 𝒞_0_, meaning both *f*_0_ and *f* can simply be a random hash function.

From the construction of the Miniception, we can recover 𝒞_0_ from 𝒪_0_. In most cases, this means we only need to store a small minimizer to implement the Miniception on the fly. In comparison, existing methods with precomputed universal hitting sets requires storage of the whole set.

Implementation of minimizers usually use monotone queues for linear time complexity, under the same assumption. The monotone queue is a special kind of deque that ensures the item in the queue are both in nondecreasing order and are inserted recent enough. Such data structure allows solving the problem of finding minimum element in every *T* -long window in input linear time, which is exactly the problem of identifying picked *k*-mers for a given sequence, as the picked locations are simply locations that are minimal in a *w*-long window. We will provide pseudocode for its implementation later this section.

As the Miniception consists of two minimizers, we use two monotone queues, one implementing the seed minimizer and one implementing the actual minimizer which identifies picked locations. As we only care whether a context is charged for the seed minimizer, the first monotone queue has expiry time *w*_0_ + 1. The second monotone queue is implemented as in a normal minimizer.

We provide a pseudocode for the algorithm below. For simplicity, we will ignore the warmup part (before the first full window). We provide a reference implementation written in Python in our github repo. We start with the monotone queue, which as described can be considered as a deque with an extra parameter *T* (expiry time) and additional operations.

The monotone queue achieves *O*(*k*) amortized time complexity (recall each item in the queue is a *k*-mer), as each item will only be inserted and popped once. With the monotone queue, we can implement the Miniception. As mentioned before, we assume each *k*-mer can be held in a word.

### Algorithm 1 The Monotone Queue with Expiry Time *T*

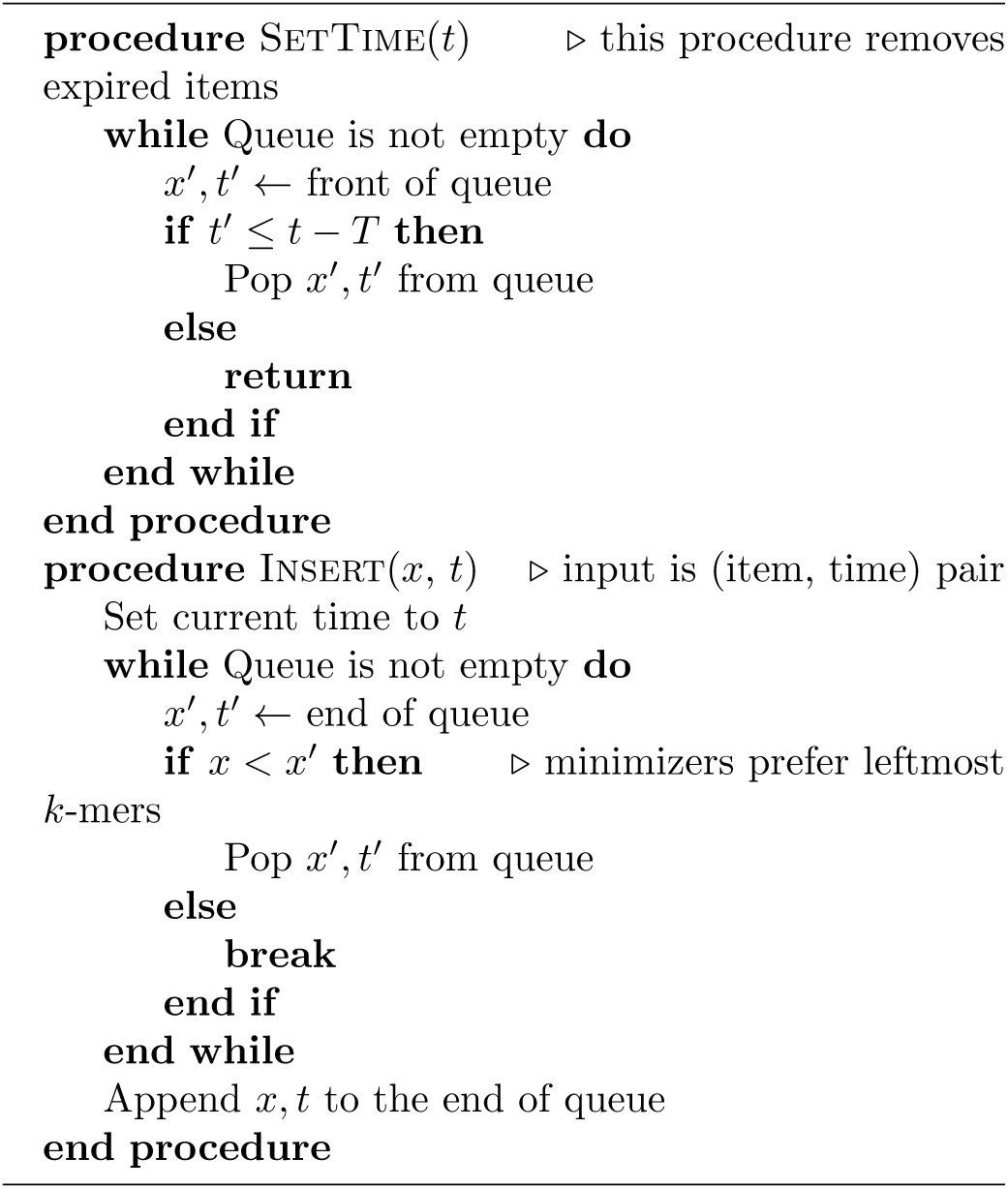

## S4 Evaluating Minimizers on Human Reference Genome

We use the identical setup as in the main text, except the contexts are now sampled from hg38 human reference genome. As observed by Roberts *et al.* (2004b,a), order the alphabet by reverse frequency may improve performance of the lexicographic minimizer. For this reason, we use the order *C* < *G* < *A* < *T* for all minimizers. We also consider two strategies for constructing a compatible minimizer given the UHS from PASHA. The authors suggested a lexicographic order within the UHS, but we also implement a minimizer where the order within the UHS is randomized, like the Miniception. Refer to the main text for more details.

### Algorithm 2 The Miniception

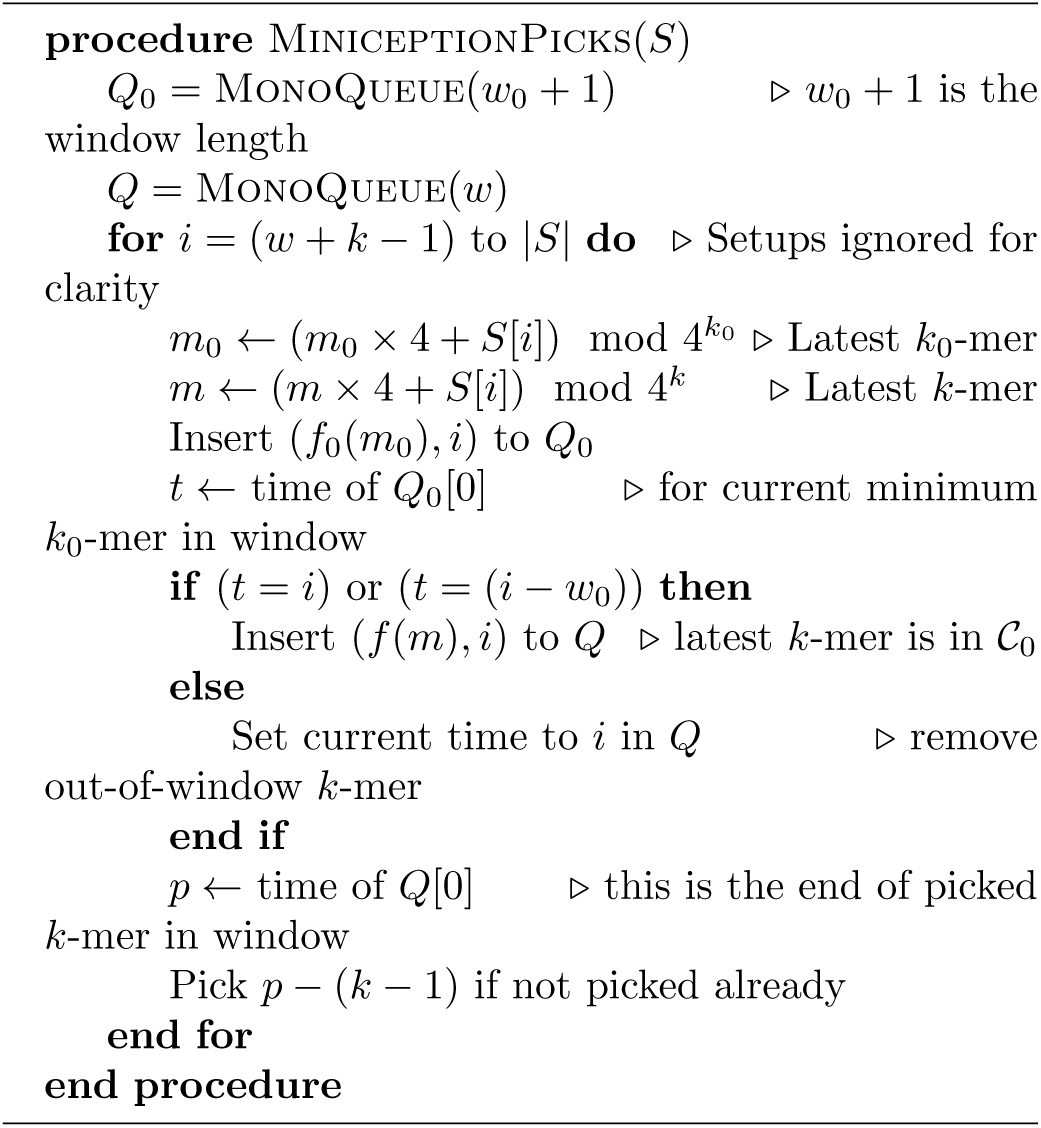

**Figure 8:**
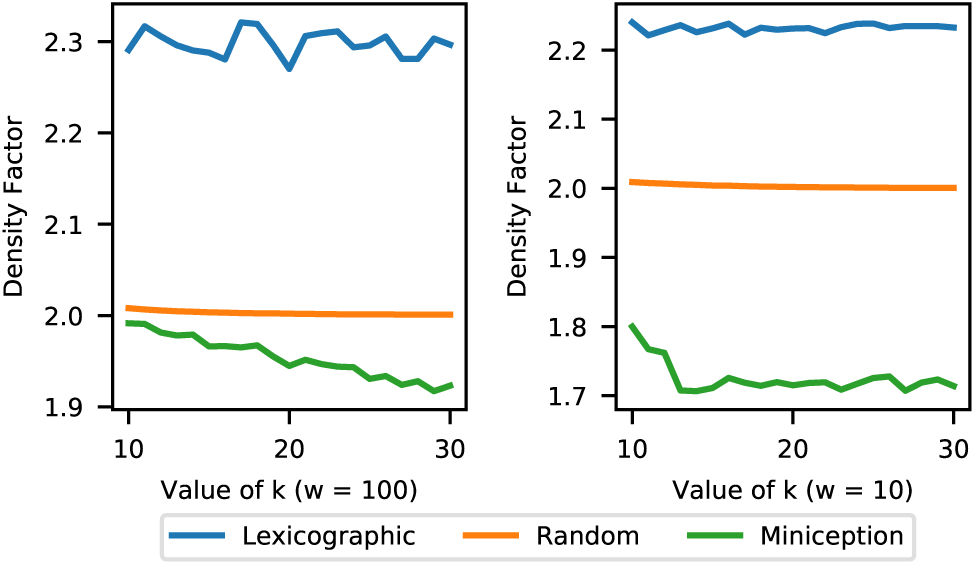
Comparing density of the Miniception against lexicographic and random minimizers. We experiment with w = 10 (left half) and w = 100 (right half).

**Figure 9:**
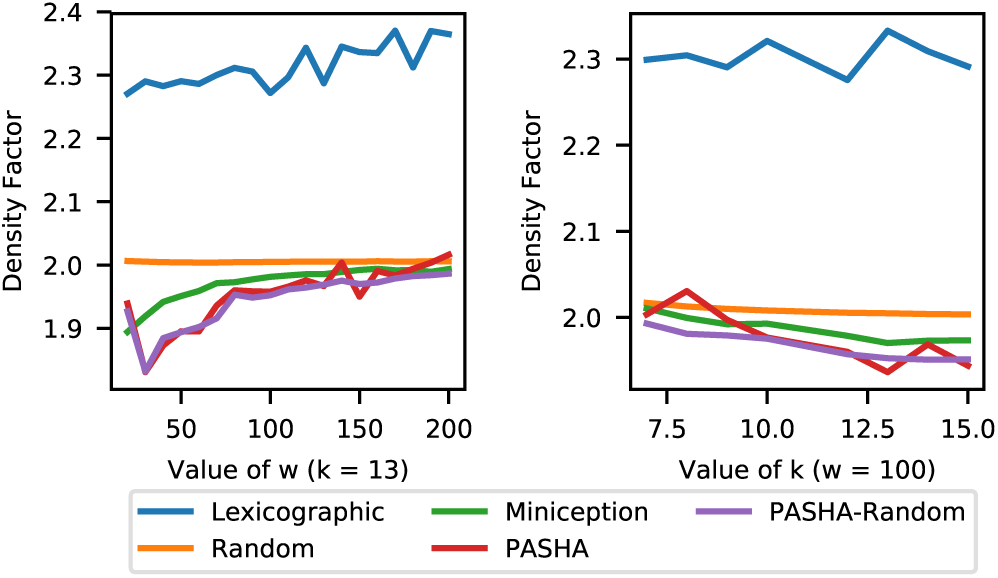
Density factor comparison, now also with minimizers derived from PASHA outputs. Left half: experiments with fixed k = 13 and varying w. Right half: experiments with fixed w = 100 and varying k.

